# TUB-010, a novel anti-CD30 antibody-drug conjugate based on Tub-tag technology, widens the therapeutic window by reducing toxicity while maintaining high efficacy

**DOI:** 10.1101/2025.01.15.633119

**Authors:** Marcus Gerlach, Saskia Schmitt, Philipp Cyprys, Marc-André Kasper, Isabelle Mai, Magdalena Klanova, Andreas Maiser, Heinrich Leonhardt, Christian P. R. Hackenberger, Guenter R. Fingerle-Rowson, Annette M. Vogl, Dominik Schumacher, Jonas Helma

**Affiliations:** Tubulis GmbH, Am Klopferspitz 19a, 82152 Planegg-Martinsried, Germany; Institute of Pathological Physiology, First Faculty of Medicine, Charles University, Prague, Czech Republic; First Department of Medicine, Department of Hematology, Charles University General Hospital, Prague, Czech Republic; Faculty of Biology, Human Biology and Bioimaging, LMU Munich, Munich, Germany; Chemical Biology, Leibniz-Forschungsinstitut für Molekulare Pharmakologie, Campus Berlin, Berlin, Germany

## Abstract

TUB-010 is a next-generation antibody-drug conjugate (ADC) targeting CD30, which is expressed on various hematopoietic malignancies such as Hodgkin lymphoma and anaplastic large cell lymphoma. Patients with refractory and relapsed CD30-positive cancers often lack effective and tolerable therapy options. Among the therapeutic options for these patients is brentuximab vedotin (Adcetris), a monomethyl auristatin E (MMAE)-delivering anti-CD30 ADC with a mean drug-to antibody ratio (DAR) of 4. Adcetris exhibits a high response rate at the cost of significant toxicities, among which neutropenia and peripheral neuropathy are the most prevalent adverse events, which are likely driven by the payload MMAE and instability of the maleimide conjugation chemistry.

TUB-010 uses the same antibody brentuximab and payload MMAE as Adcetris, but instead of maleimide chemistry, TUB-010 is based on the Tub-tag conjugation strategy, which enables the generation of a homogenous and site-specific DAR 2 ADC with unique biophysical properties. This new technology stably attaches MMAE to the hydrophilic Tub-tag peptides on the light chains via chemoenzymatic conjugation using the enzyme tubulin tyrosine ligase.

TUB-010 demonstrates similar binding affinity, internalization and lysosomal release characteristics as Adcetris in CD30-positive cells. When normalized to the MMAE concentration, TUB-010 shows comparable *in vitro* cytotoxic efficacy as well as similar bystander activity compared to Adcetris on established cancer cell lines. Importantly, TUB-010 exhibits higher stability with neglectable premature deconjugation in circulation and reduced high molecular weight species formation as well as lower non-specific cytotoxicity on target-negative cells compared to Adcetris. As a consequence, TUB-010 induces superior tumor control compared to Adcetris when dosed at equal MMAE concentrations *in vivo* and also shows lower toxicity and higher tolerability in rodents and non-human primates.

Taken together, TUB-010 is a novel and potential best-in-class anti-CD30 ADC with improved biophysical properties designed to deliver the cytotoxic payload with higher precision and with a wider therapeutic window than Adcetris using Tub-tag conjugation technology. Therefore, TUB-010 may increase the clinical benefit of ADC therapies for patients with CD30-positive malignancies.

## Introduction

CD30-positive lymphomas represent a heterogeneous group of hematologic neoplasms arising from malignant transformation of normal B or T lymphocytes at various stages of their development. CD30 (also known as TNFRSF8) is a transmembrane glycoprotein belonging to the tumor necrosis factor (TNF) receptor alpha superfamily (1). CD30 is highly and uniformly expressed by malignant cells of classic Hodgkin lymphoma (cHL) and anaplastic large cell lymphoma (ALCL), for both of which CD30 expression is a hallmark of diagnosis, as well as to various levels by a substantial proportion of non-ALCL peripheral T cell lymphomas (PTCL) or primary cutaneous T cell lymphomas (CTCL) and by certain subtypes of B cell non-Hodgkin lymphomas (2-6). CD30 expression in healthy tissues is restricted mainly to subsets of activated T and B cells (7-9). Signaling through CD30 can have pleiotropic effects on cell proliferation and survival depending on the cell type and cell state and is mediated mainly by nuclear factor kappa B and mitogen-activated protein kinase/extracellular signal-regulated kinase pathways (10-13). In addition, CD30 antigen rapidly internalizes into the cell upon antibody binding, making it an ideal target for selective drug delivery (14).

Brentuximab vedotin (Adcetris) is a CD30-directed antibody-drug conjugate (ADC) composed of the humanized IgG1 monoclonal antibody brentuximab (cAC10) and approximately four linker-payload moieties consisting of a thiol-reactive maleimide, a caproyl spacer, the protease-cleavable linker valine-citrulline-*p*-aminobenzyloxycarbonyl (VC-PAB) and the microtubule-disrupting agent monomethyl auristatin E (MMAE). The maleimide attachment group covalently binds the linker-payload construct (MC-VC-PAB-MMAE) to cysteine residues of the cAC10 antibody (15).

The maleimide-based technology used for conjugation of MC-VC-PAB-MMAE to the monoclonal antibody cAC10 results in an ADC with drug-to-antibody ratio (DAR) ranging between 0 to 8, with an average DAR (DAR_av_) of 4. In addition, maleimide-conjugated linker-payload is prone to undesirable transfer of linker-payload to serum proteins during blood circulation, mediated by retro-Michael addition (16).

Adcetris has been approved for several indications in cHL and systemic ALCL (previously untreated and relapsed/refractory [R/R] setting), CD30+ non-ALCL PTCL (previously untreated), and CD30+ CTCL following prior systemic treatment based on the results of several phase II-III clinical trials (17-23). The long clinical experience with Adcetris has brought important insights into the safety profile of the drug. Neurotoxicity and hematological toxicity, especially neutropenia, represent off-target toxicities that belong to the most frequent clinically relevant adverse events reported across multiple clinical trials with Adcetris. Peripheral neuropathy represents a clinical challenge, as it occurs with high frequency (ranging between 40% to 56%) and represents the primary cause of dose delay, dose modifications or premature discontinuation of Adcetris (18, 20, 21, 23).

In this study we describe the preclinical development of TUB-010, a next generation CD30-targeting ADC. In contrast to Adcetris, TUB-010 uses Tub-tag conjugation to stably attach the linker-payload (hydroxylamine[HA]-VC-PAB-MMAE) to the light chains of the monoclonal antibody cAC10 (24, 25). The Tub-tag is a 14 amino acid peptide derived from the C-terminus of α-tubulin and is expressed at the C termini of the light chains of cAC10. The Tub-tag technology produces a homogenous (DAR 2), highly stable anti-CD30 ADC with improved biophysical properties resulting in high efficacy and better tolerability compared to Adcetris in preclinical *in vivo* studies.

## Materials and Methods

### ADC conjugation

Chemoenzymatic conjugation of linker-payload (HA-VC-PAB-MMAE) to the monoclonal antibody is accomplished by Tub-tag technology as previously described (24, 25). In the first step, the terminal glutamic acid of Tub-tag is enzymatically functionalized by attachment of 3-formyl-*L*-tyrosine using tubulin tyrosine ligase (TTL). After removal of excess TTL, 3-formyl-*L*-tyrosine and buffer exchange, the linker-payload is covalently attached by bioorthogonal oxime-ligation. After completion of the conjugation reaction excess of HA-VC-PAB-MMAE was for example removed by ultrafiltration/diafiltration.

Adcetris used as a reference was purchased from Takeda (Japan).

### Evaluation of HMWS formation with HPLC-SEC

High performance lipid chromatography (HPLC)-size exclusion chromatography (SEC) was used for the quantification of high molecular weight species (HMWS) formed during storage at elevated temperatures (e.g. 40°C) and freeze-thaw cycles (up to 5 cycles) according to supplementary materials. At the start of each study, TUB-010 and Adcetris were adjusted to a similar protein concentration and the ADCs were transferred into the same formulation. Sterile filtration was carried out prior to storage.

### Evaluation of serum stability via LC-MS

Human serum samples were spiked with either TUB-010 or Adcetris (0.3 mg/mL) and stability was assessed by pulldowns followed by LC-MS analysis and calculation of the DAR_av_ from the obtained spectra.

Pulldowns were performed with NHS-activated magnetic beads (Thermo Fisher Scientific, USA) that were coupled to a commercial anti-cAC10 antibody (R&D Systems, USA; Cat# MAB9584, RRID:AB_3659482). Coupling was performed according to the manufacturer’s instructions, using 0.33 µg of antibody per 1 µL of beads. For pulldowns, 100 µL of spiked serum sample was diluted with 200 µL of PBS. 40 µL of anti-cAC10 beads were added, and the samples were gently rotated for 2 hours at room temperature. The beads were washed 2 times with TBS-T followed by elution with 100 mM glycine pH 2.5 for 10 minutes at 900 rpm. Eluates were neutralized with 1/10 volume of 1 M tris base pH 10.5 and buffer-exchanged into PBS using Zeba Spin gel filtration columns (Thermo Fisher Scientific, USA) followed by reduction (5 mM DTT) and deglycosylation with PNGase F (0.5 U/µL, Promega, USA) at 37°C for 2 hours. LC-MS analysis was performed as described in the supplements. DAR_av_ was calculated from the obtained spectra.

### *In vitro* cytotoxicity and bystander effect

CD30-positive cells were incubated for 4 days with increasing concentrations of TUB-010, Adcetris and cAC10 up to a maximum concentration of 3 µg/mL or with free MMAE up to 1 µM. Killing was analyzed using resazurin cell viability dye (Merck, Germany) by dividing the fluorescence of ADC-treated cells from control cells in medium. Fluorescence emission at 590 nm was measured on a microplate reader Infinite M1000 Pro (Tecan, Switzerland).

Alternatively, cell viability was analyzed by CellTiter-Glo Luminescent Cell Viability Assay (Promega, USA) according to the manufacturer’s instructions and luminescence was measured on a microplate reader Infinite M1000 Pro (Tecan, Switzerland) in flat white 96-well plates (Corning, USA).

Target-negative healthy human cells were incubated with TUB-010 and Adcetris concentrations up to 100 µg/mL for 7 days to promote unspecific toxicity followed by resazurin-based viability readout as described above.

For the supernatant transfer-based bystander assay, CD30-positive cells were incubated with increasing concentrations of TUB-010, Adcetris and cAC10-MMAF up to a maximum concentration of 3 µg/mL. cAC10-MMAF was conjugated using the proprietory P5 technology to a DAR of 8 (26-28) and used as a negative control. After 4 days, cell supernatant was transferred to CD30-negative HL-60 cells and incubated for another 4 days. Killing was analyzed using resazurin-based viability measurement as stated before.

For the co-culture-based bystander assay, CD30-positive cells (Karpas-299 and L-540) and CD30-negative HL-60 cells were incubated at an 8:1 ratio with increasing concentrations of TUB-010, Adcetris and cAC10-MMAF up to a maximum concentration of 3 µg/mL. After 3 days, fresh ADC solutions were added to the co-cultures to feed the cells. After another 3 days incubation time, cocultures were stained by αCD25-FITC (BioLegend, UK; Cat# 356105, RRID:AB_2561862), αCD33-APC (BioLegend, UK; Cat# 303408, RRID:AB_314352) and Fixable Aqua Dead Cell Stain Kit (Thermo Fisher Scientific, USA). As a readout for the bystander effect, the percentage of dead CD30-positive (Karpas-299, L-540, CD25-positive) and -negative (HL-60, CD33-positive) was determined by flow cytometry.

### Mouse xenograft experiments

*In vivo* efficacy studies were conducted at Experimental Pharmacology & Oncology Berlin-Buch (EPO), Germany and Charles River Discovery Research Services, Germany, in accordance with the local animal welfare laws and approved by local authorities. Xenografts were initiated with 1×10^7^ Karpas-299 cells per mouse subcutaneously injected in the right flank of 6–8-week-old female CB17-severe combined immunodeficiency (SCID) mice (Janvier Labs and Charles River; RRID:IMSR_RJ:CB17-SCID) on day 0. Animals were randomized into treatment and control groups with indicated numbers of experimental animals per group when tumor volumes reached 100–200 mm^3^. Test articles, TUB-010 and Adcetris, or vehicle control were administered intravenously (i.v.) via tail vein injection. Mice were dosed at doses and frequencies as indicated in the experimental description. Tumors and body weights were measured twice to three times per week. Study endpoints were tumor volume (TV) of >1500 mm^3^, distress of the animals or day 57, 54, or 80 for the three studies, whichever came first.

### Tissue cross-reactivity study

Tissue cross-reactivity (TCR) studies were conducted at Labcorp Early Development Laboratories Ltd. Histologically normal frozen human tissues, collected from donors cryo-sectioned or prepared as smears, were used in this study. CD30 protein spots served as positive control. The tissues were stained immunohistochemically with vimentin, cytokeratin and von Willebrand factor to assess the viability. To facilitate immunohistochemical detection of TUB-010 and control article, the test articles were biotinylated (ImmunoProbe Biotinylation Kit, Merck, Germany). The detection was carried out using a rabbit anti-biotin antibody (Abcam, UK; Cat# ab53494, RRID:AB_867860) in combination with an anti-rabbit HRP secondary antibody (Agilent, USA; Cat# K4003, RRID:AB_2630375).

### Toxico-(TK) and pharmacokinetics (PK) studies in non-human primates

PK/TK studies in naïve male and female cynomolgus monkeys were performed at Labcorp Early Development Laboratories Ltd. The studies and all procedures followed United Kingdom National Law, in particular the Animals (Scientific Procedures) Act 1986.

Groups of cynomolgus monkeys (n = 2, female) were given an i.v. infusion of vehicle control or 6, 12 or 15 mg/kg of TUB-010 once every 3 weeks (Q3W) for a total of four cycles (6 mg/kg group) or two cycles (12 and 15 mg/kg of TUB-010). Multiple parameters were monitored during the study including mortality, clinical and post-dose observations, body weight and food consumption. Blood sample evaluation for PK/TK, hematology, coagulation and clinical chemistry and urinalyses were performed pre-dose and at selected time points throughout the study. Necropsy was conducted on day 71 of the study (6 mg/kggroup) or on day 43 (12 and 15 mg/kgof TUB-010) including macroscopic examinations, organ weights tissue preservation and fixation as well as histology, microscopic evaluation.

For PK analysis, blood was sampled at pre-dose and after 0.5, 1, 4, 24, 48, 72, 168, 336 and 504 h. Total antibody (mAb) and intact ADC were analyzed by ELISA methods as described in the Supplementary Information.

## Results

### TUB-010 is a homogenous DAR2 anti-CD30-MMAE ADC based on Tub-tag technology

The site-specific chemoenzymatic Tub-tag conjugation technique, on which TUB-010 is based, is derived from the post-translational addition of tyrosine to the C-terminal sequence of mainly α-tubulin within the tubulin hetero-dimer catalyzed by the enzyme tubulin tyrosine ligase (TTL) (Fig. 1A) (24, 25, 29). Within cells, the unstructured C-terminal α-tubulin tail protrudes to the outside of the assembled microtubule lattice – one of the major constituents of the eukaryotic cytoskeleton – thereby being readily accessible and subjected to various post-translational modifications regulating dynamics of the microtubules, as well as their organization and interaction with other cellular components. Employed as a protein tag, recombinantly expressed at the C-terminus of proteins such as antibody chains, the highly negatively charged 14 amino acid sequence (VDSVEGEGEEEGEE) provides a favorable hydrophilic microenvironment that can be exploited for site-specific conjugation of highly hydrophobic chemical moieties such as linker-payload structures employed for ADCs. In agreement with this property, the Tub-tag sequences expressed at the C-termini of light chains of monoclonal antibodies, here shown for a monoclonal IgG1 antibody with the sequence of brentuximab (called cAC10 LC Tub throughout the manuscript), resulted in significantly reduced retention times during HPLC-HIC analysis, compared to the unmodified antibody (called cAC10) indicative of increased hydrophilicity (Fig. 1B).

**Figure 1.**
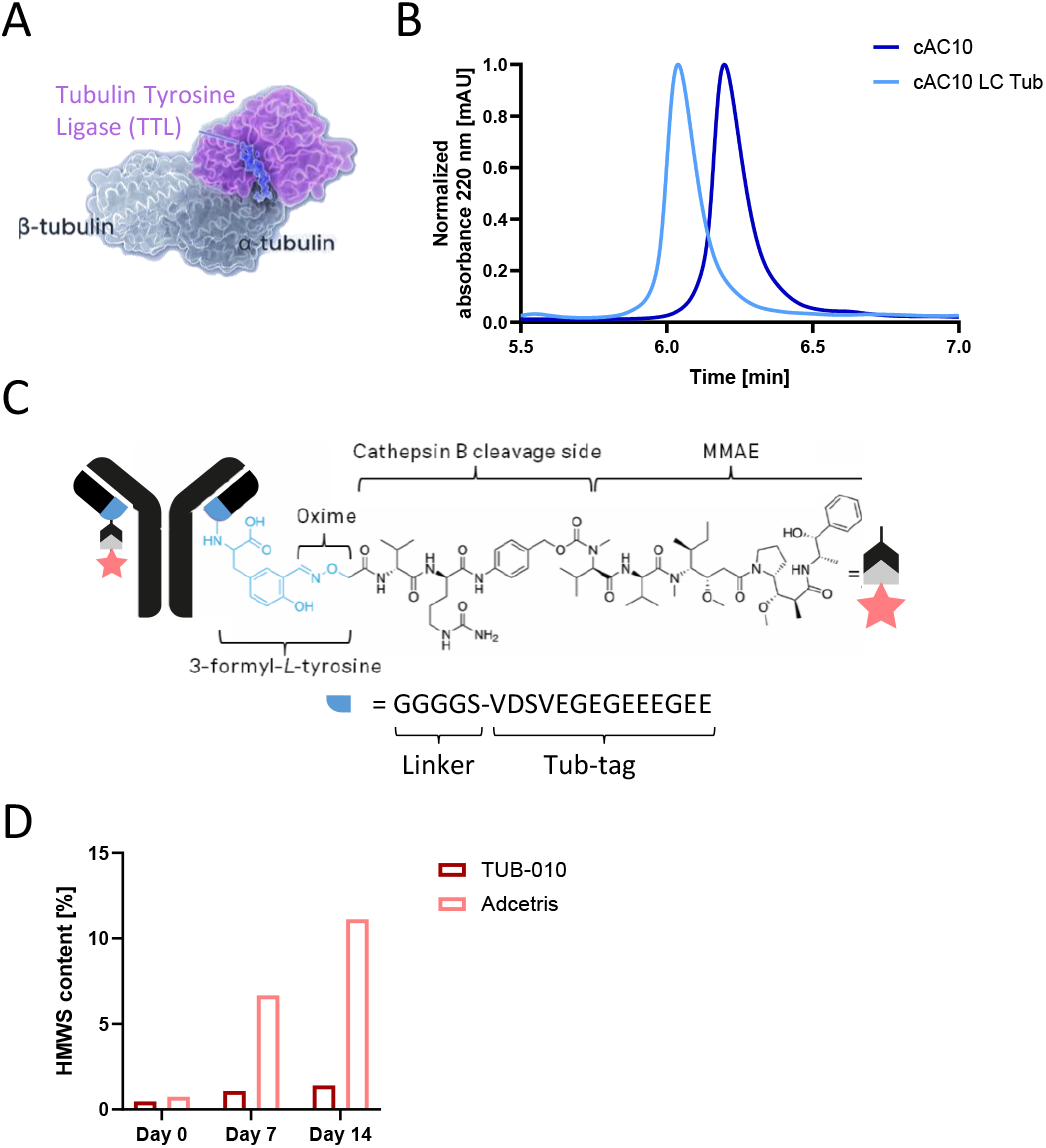
TUB-010 is a homogenous ADC based on Tub-tag technology and shows reduced HMWS formation at elevated temperatures. **(A)** Tub-tag conjugation technology derived from microtubule biology. Depicted is tubulin-tyrosine ligase (in pink) catalyzing the enzymatic addition of tyrosine to the C-terminal sequence of alpha-tubulin (in blue) within the tubulin hetero-dimer consisting of an α-tubulin (dark grey) and β-tubulin (light grey) subunit. **(B)** HPLC-HIC analysis of unmodified antibody (cAC10) and LC Tub-tagged cAC10 (cAC10 LC Tub). **(C)** Schematic drawing and composition of TUB-010, a homogenous DAR 2 anti-CD30-MMAE Tub-tag ADC. **(D)** HMWS formation indicating aggregation measured by HPLC-SEC after incubation of TUB-010 and Adcetris for up to 14 days at 40°C.

TUB-010 is a next generation CD30-targeting ADC, consisting of cAC10 LC Tub conjugated to the payload MMAE, a microtubule-targeting cytotoxin, via the cleavable valine-citrulline *p*-aminobenzylcarbamate (VC-PAB) linker using Tub-tag conjugation. In the first conjugation step, TTL adds 3-formyl-*L*-tyrosine to the C-termini of the Tub-tag sequences, followed by the second step of linker-payload (HA-VC-PAB-MMAE) addition via chemoselective oxime ligation resulting in a homogenous ADC (Fig. 1C, Supplementary Fig. S1). Of note, TUB-010 showed > 5-fold reduced HMWS formation under stress conditions compared to Adcetris (Fig. 1D), attributable to improved biophysical properties. This is on the one side mediated by the hydrophilic Tub-tag sequences, which counterbalance the hydrophobic linker-payload structure, and on the other side by the site-specific homogenous conjugation compared to the maleimide conjugation technique used in Adcetris which results in an aggregation-prone heterogenous DAR 0-8 ADC product. cAC10 LC Tub and unmodified cAC10 exhibited highly similar binding and *K*_D_ values in enzyme-linked immunosorbent assay (ELISA)-based and cellular binding assays, indicating that the addition of the Tub-tag sequence does not influence antigen binding properties of the antibody (Supplementary Fig. S2A and S2B). Moreover, cAC10 LC Tub antibody and TUB-010 ADC showed similar internalization behavior on CD30-positive cells using pHrodo conjugates (Supplementary Fig. S2C). Lysosomal accumulation of TUB-010 was further confirmed by co-localization with the lysosomal marker LAMP1 using super-resolution microscopy (Supplementary Fig. S2D).

### TUB-010 shows minor payload loss and excellent *ex vivo* and *in vivo* serum stability

*Ex vivo* stability and loss of linker-payload of TUB-010 and Adcetris was assessed by incubation of the ADCs in sera from different species followed by immunoprecipitation and analysis of the DAR_av_ by LC-MS. Adcetris showed to be highly unstable with major linker-payload loss (75.8% - 78.3%) in all three sera after 7 days of incubation at 37°C. In contrast, TUB-010 is remarkably stable and even after a prolonged incubation period of 21 days only minor loss of linker-payload (19.5% - 20.5%) was observed (Figure 2A). *In vivo*, a high percentage of non-specific transfer of linker-payload to serum proteins was confirmed for Adcetris that is mediated by retro-Michael addition triggered from the maleimide conjugation used in Adcetris, leading to a reduced linker-payload content present on the ADC (30, 31). In contrast, TUB-010 only showed neglectable linker-payload transfer to serum proteins in circulation (Fig. 2B). In line with this finding, when pre-incubated in serum, Adcetris showed reduced cytotoxicity on CD30-positive cells, whereas pre-incubated TUB-010 remained equally efficient. No cytotoxicity was seen on target-negative cells (Supplementary Fig. S3A-B). Overall, TUB-010 exhibited excellent *in vivo* stability in cynomolgus monkeys and rats, with super-imposable total antibody and intact ADC curves specifically for early time points, and almost antibody-like *in vivo* PK with no evidence of increased clearance (Fig. 2C, Supplementary Fig. S3C-D). In addition, TUB-010 showed lower free MMAE levels compared to Adcetris (Supplementary Fig. S3E). In summary, these data sets confirm the high stability and only minimal premature extracellular payload loss of TUB-010.

**Figure 2.**
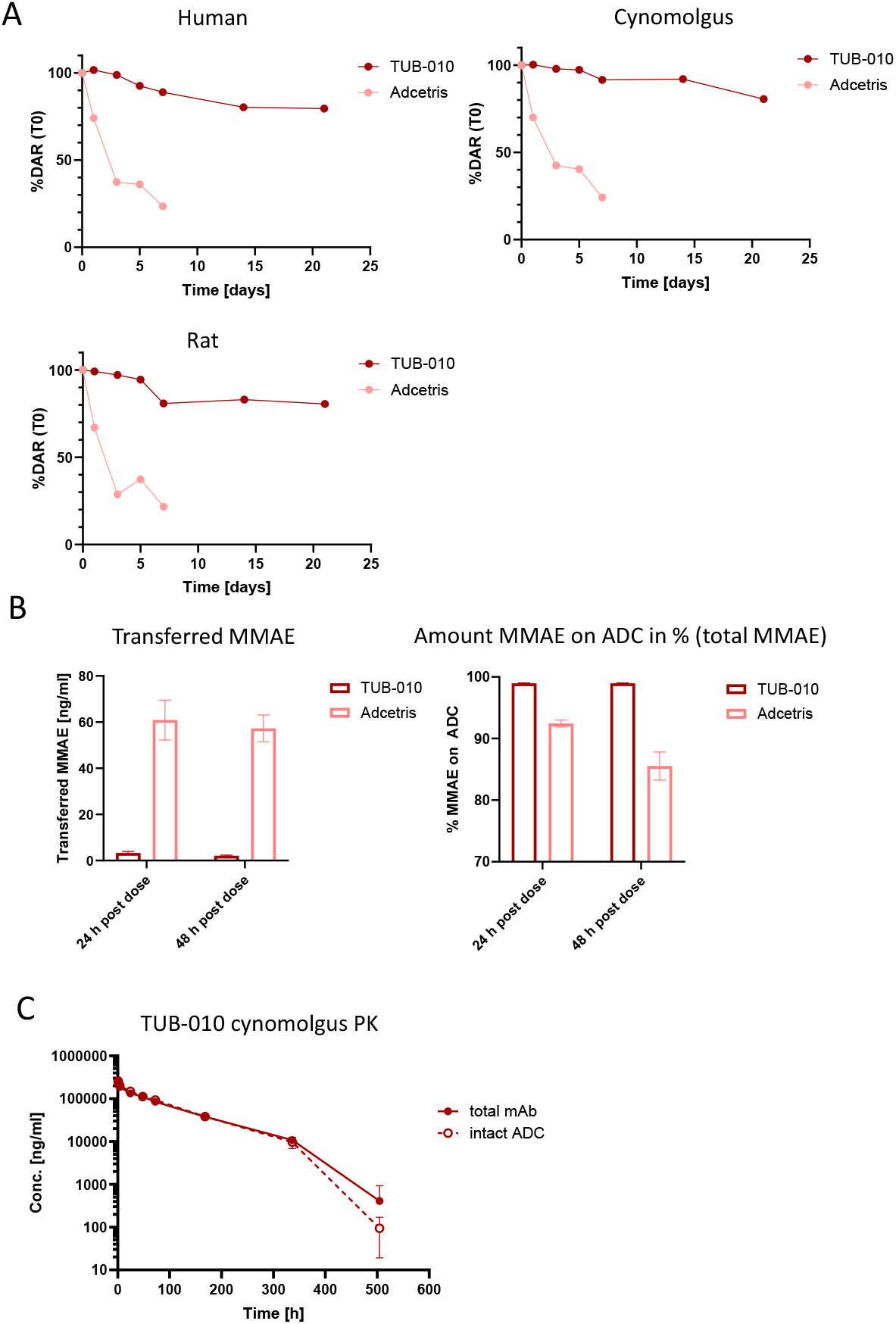
Stability of TUB-010 and Adcetri. **(A)** *Ex vivo* serum stability of Adcetris and TUB-010. ADCs were incubated for up to 21 days in human, cynomolgus or rat serum at 37°C. ADCs were re-isolated by immunoprecipitation and the DAR_av_ was analyzed by LC-MS. Shown is the % of DAR compared to time point 0 (T0). **(B)** Pharmacokinetics analysis of TUB-010 and Adcetris in rats. MMAE transferred to serum proteins and amount of MMAE on ADC was measured in serum samples by LC-MS. Graph shows mean ± SD, n = 2. **(C)** Pharmacokinetic analysis of TUB-010 in cynomolgus monkey. Total antibody and intact ADC were measured by ELISA after indicated sampling times in serum samples from cynomolgus monkeys dosed with 12 mg/kg TUB-010 i.v. Total ADC was captured using an anti-idiotype brentuximab antibody, intact ADC by an anti-MMAE antibody. Data points represent mean ± SD, n = 2.

### TUB-010 shows efficient *in vitro* cytotoxicity and bystander activity and improved *in vivo* tumor control compared to Adcetris when dosed at equal MMAE concentrations

TUB-010 showed *in vitro* cytotoxicity on ALK-positive ALCL, cHL, PTCL, ALK-negative ALCL, and CTCL cancer cell lines with EC_50_ values ranging from 0.02 and 0.23 nM in most cell lines correlating with CD30 surface expression and sensitivity of cell lines towards free MMAE (Fig. 3A and Supplementary Fig. S4A). The HL cell line L-428 exhibits overexpression of MDR1 and therefore resistance towards TUB-010 with an EC50 of only 921.7 ng/mL (in-house data and (32, 33)). When normalized to the MMAE concentration, TUB-010 revealed similar *in vitro* cytotoxic efficacy as well as similar bystander activity compared to Adcetris on *in vitro* cancer cell lines from different indications (Supplementary Fig. S4B–C, Supplementary Fig. S5). Moreover, TUB-010 and Adcetris showed highly similar antibody-dependent cellular cytotoxicity (ADCC), complement-dependent cytotoxicity (CDC), and antibody-dependent cellular phagocytosis (ADCP). Both agents were capable of inducing ADCP, but did not exhibit CDC and ADCC activity (Supplementary Fig. S6A–E). In addition, TUB-010 was able to induce dendritic cell (DC) maturation to a similar extent as free MMAE, most likely via a combination of induction of immunogenic cell death of cancer cells as measured by the release of damage-associated molecular patterns (DAMPs) as surrogate markers and direct immune stimulatory effect of released MMAE via the bystander effect as described previously (Supplementary Fig. S6F and S6G) (34). The excellent *in vitro* cytotoxicity and high serum stability of TUB-010 translated to a similar tumor regression and survival of TUB-010-compared to Adcetris-treated mice in a repeated dose study in the Karpas-299 xenograft model when both ADCs are dosed at the same protein concentration with TUB-010 only delivering half the amount of MMAE (TUB-010 being a DAR 2 ADC, Adcetris being a DAR 4 ADC) (Fig. 3B). TUB-010 was found active in dose response with 7/10 complete regressions (CRs) at the minimal effective dose of 1 mg/kg and 9/10 CRs at 1.5 mg/kg. Interestingly, when dosed at equal MMAE concentrations *in vivo* (10 µg/kg), TUB-010 improved tumor control with 70% CRs for TUB-010 versus 40% for Adcetris, and prolonged survival in the Karpas-299 xenograft model compared to Adcetris (Fig. 3C).

**Figure 3.**
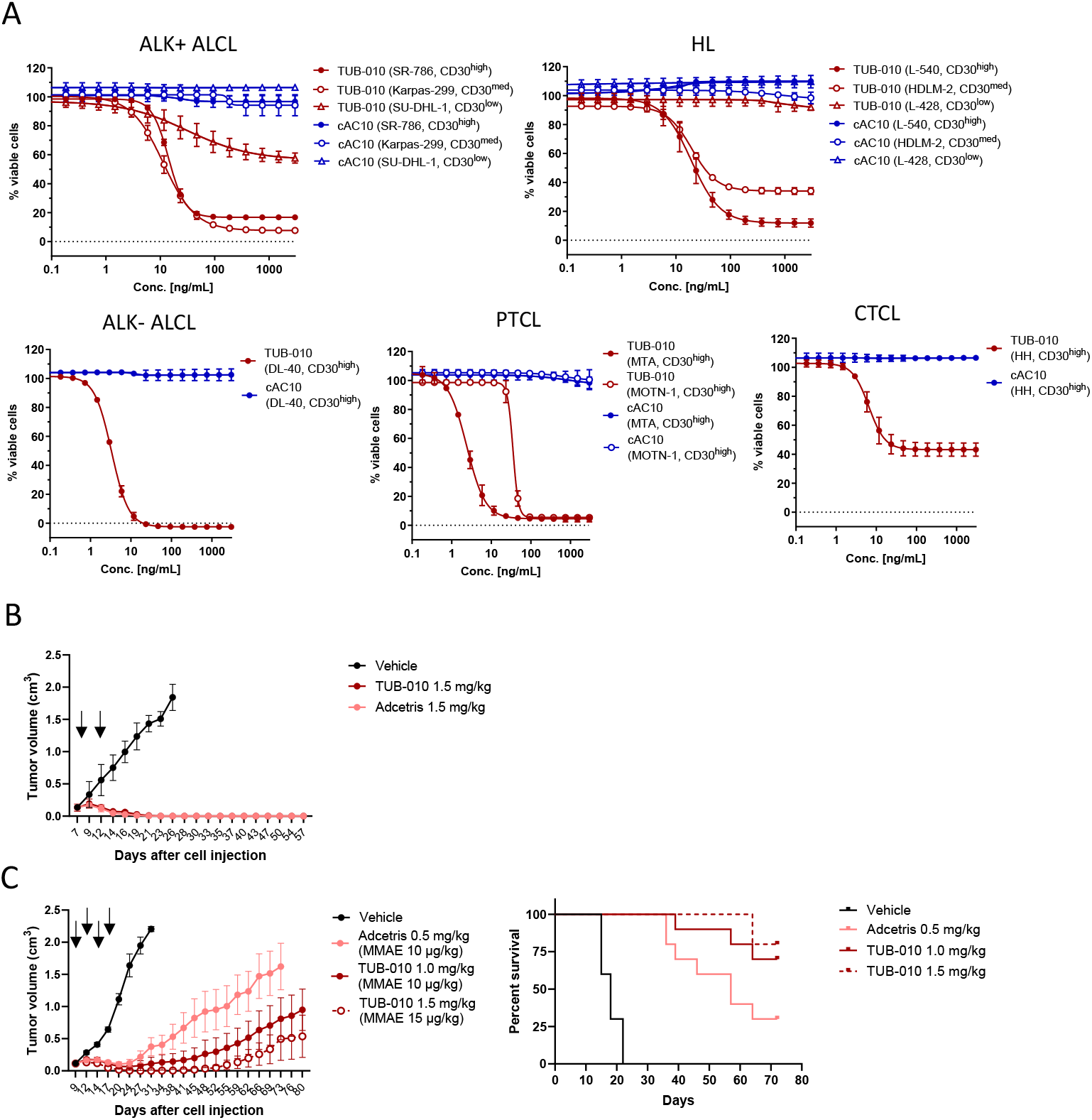
TUB-010 shows highly efficacious *in vitro* cytotoxicity and is more efficacious than Adcetris when dosed at similar MMAE concentrations in xenograft model. **(A)** *In vitro* cytotoxicity of TUB-010 in cell lines from ALK-positive ALCL, HL, PTCL, CTCL and ALK-negative ALCL cancers expressing varying CD30 levels. Cells were incubated for 4 days with increasing concentrations of TUB-010 or cAC10 and viability was analyzed using a resazurin-based readout. Data points show mean ± SD. Shown are representative killing curves of n = 2-3 individual experiments. **(B-C)** *In vivo* anti-tumor efficacy of TUB-010 vs Adcetris in Karpas-299 xenograft model. Tumor-bearing mice were intravenously administered with TUB-010 or Adcetris at the indicated concentrations when tumor volumes reached 100-150 mm^3^ at day 7 and 10 **(B)** and at day 9, 12, 14, and 17 **(C)** after tumor cell injection. Graphs show mean tumor volume ± SEM and Kaplan Meyer survival plots, n = 8 (B), n = 10 (C).

### Tub-tag technology results in reduced non-specific uptake and cytotoxicity in target-negative cells compared to Adcetris

Of note, TUB-010 showed reduced non-specific cellular uptake and cytotoxicity in a panel of CD30-negative cells, including primary human cells that are established *in vitro* toxicity models (Fig. 4A and 4B). When normalized to the MMAE concentration, TUB-010 lowered the cellular toxicity mediated by target-independent non-specific uptake by factor 2-to >10-fold in several human primary cells within the clinically relevant therapeutic range (5-50 µg/mL) compared to Adcetris (Fig. 4C) (35). We hypothesized that the increased hydrophilicity of the overall ADC introduced by the Tub-tag sequences and the reduced aggregation behavior mediate the reduced nonspecific cellular uptake and consecutively the reduced non-specific cellular toxicity. Also, the negatively charged Tub-Tag might lead to repulsion on the negatively charged membrane.

**Figure 4.**
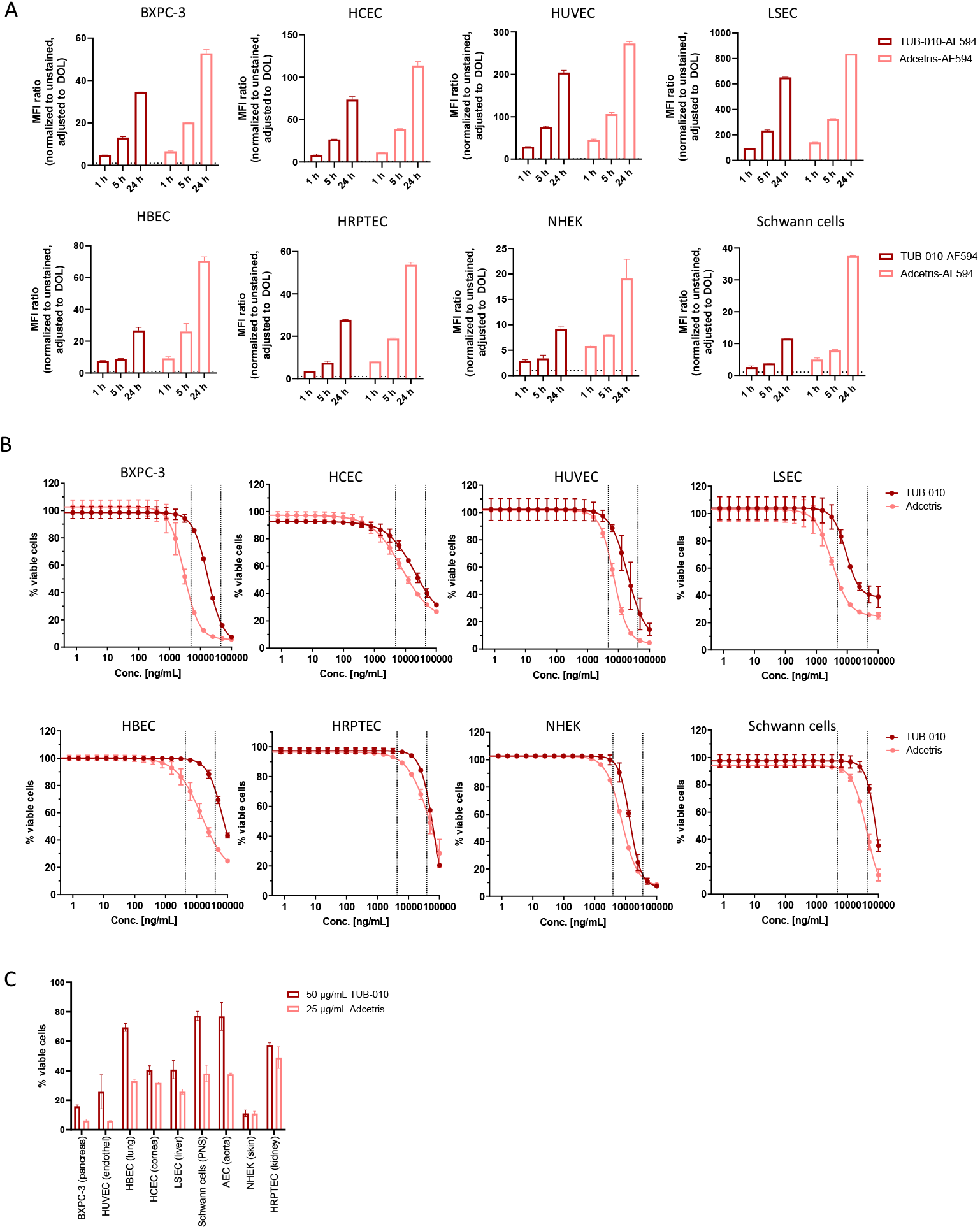
TUB-010 induces lower non-specific internalization and cytotoxicity of target-negative cells compared to Adcetris. **(A)** Non-specific internalization of TUB-010 and Adcetris in CD30-negative cells. Indicated human primary cells and cell lines were incubated for indicated time points with AF594-labeled TUB-010 and Adcetris and analyzed by flow cytometry. Graphs show means of n = 2 ± SD. **(B)** Indicated CD30-negative cells were incubated for 7 days with increasing concentrations of TUB-010 and Adcetris up to 100 µg/mL. Cytotoxicity was analyzed using a resazurin-based cell viability readout. Data points for Adcetris were normalized to the MMAE amount of TUB-010 (in ng/mL). Dotted lines indicate the range of popPK model predicted Adcetris concentrations (5-50 µg/mL) in patients with hematological malignancies (35). **(C)** Off-target cytotoxicity on target-negative cells at selected clinically relevant concentrations. All graphs show means of n = 2 ± SD. Abbreviations, BXPC-3: pancreatic cancer cell line, HCEC: human corneal endothelial cells, HUVEC: human umbilical vein endothelial cells, LSEC: liver sinusoidal endothelial cells, THLE-3: transformed human liver epithelial cells, HBEC: human bronchial epithelial cells, HRPTEC: human renal proximal tubular epithelial cells, NHEK: normal human epidermal keratinocytes, Schwann cells: human glia cells of the peripheral nervous system, iAEC: immortalized aortic endothelial cells.

### TUB-010 shows lower toxicity and higher tolerability in rodents and non-human primates compared to Adcetris indicating a higher therapeutic window

The binding of TUB-010 to 40 human tissues was analyzed in a tissue cross-reactivity study, revealing a highly similar and highly restricted binding pattern of TUB-010 compared to published Adcetris data (36). None of the tested normal human tissues showed binding of TUB-010.

The safety and toxicology profile of TUB-010 was evaluated in repeated dose studies in rats (non-cross-reactive species) and cynomolgus monkeys (cross-reactive species). In the repeated dose rat study, animals were dosed i.v. 4 times Q1W with either 10 mg/kg TUB-010 or Adcetris. As expected from published data (36), myelotoxicity was the primary test item-related toxicity with Adcetris. At the same dose and dosing schedule, these hematological toxicities were markedly attenuated with TUB-010. In cynomolgus monkey toxicity and toxicokinetic studies, TUB-010 was administered 2 or 4 times Q3W in a dose range of 6-15 mg/kg. In these studies, TUB-010 showed reduced hematological toxicity compared to published Adcetris data (36), with neutropenia being the most severe side effect at high doses. No major changes in clinical chemistry parameters and no signs of anatomical or functional tissue damage were detectable. The maximum tolerated dose (MTD) was not reached, the highest non severely toxic dose (HNSTD) of TUB-010 was determined at 12 mg/kg and the no observed adverse effect level (NOAEL) was observed at 6 mg/kg (Fig. 5). In summary, the safety and toxicology studies showed a ∼5-fold widened therapeutic window for TUB-010 compared to Adcetris with a MTD of 3 mg/kg (36) when dosed at the same protein concentration and a ∼2.5-fold widened therapeutic window when dosed at the same MMAE concentration. Thus, the presented data is supportive of a higher potential dosing of TUB-010 in patients in the clinic to achieve higher efficacy and improved tumor control while maintaining acceptable toxicity profiles.

**Figure 5.**
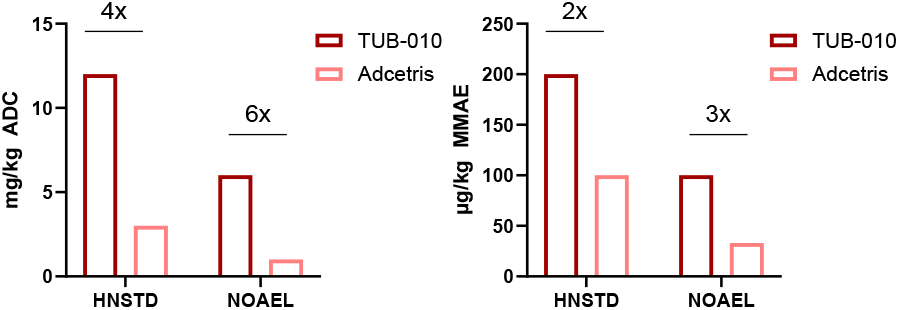
TUB-010 toxicology assessment in non-human primates supports higher tolerated dosing compared to Adcetris. Toxicology studies in non-human primates revealed a higher HNSTD and NOAEL for TUB-010 compared to Adcetris indicating a widened therapeutic window for TUB-010. HNSTD: highest non severely toxic dose, NOAEL: no observed adverse event level.

## Discussion

ADCs represent a rapidly emerging class of therapeutic agents that combine the target specificity of an antibody with the potent cytotoxic effects of a linked drug. Adcetris, a CD30-targeting ADC significantly contributed to the success of ADCs due to its proven clinical activity across multiple CD30+ hematological malignancies. To date, Adcetris in combination with chemotherapy has been approved as frontline treatment for patients with advanced cHL or ALCL, as well as other CD30-positive PTCL in the US. It is also approved as a single agent in several indications including R/R cHL, ALCL or CTCL. Adcetris uses the maleimide chemistry to conjugate the microtubule inhibitor MMAE to the anti-CD30 mAb brentuximab (cAC10) through a protease-cleavable VC-PAB linker. The high specificity of CD30 as a target combined with the potency of MMAE predestined Adcetris to become an effective drug. In contrast, the maleimide-based technology used in development of Adcetris is the source of shortcomings that limit the efficacy and negatively impact the safety of the drug.

To address these challenges, we aimed to build on the successful proven parts of Adcetris and develop a next generation CD30-targeting ADC using the novel Tub-tag conjugation technology. This approach stably attaches HA-VC-PAB-MMAE to the light chains of the monoclonal antibody cAC10 via the Tub-tag, a 14 AA peptide expressed at the C termini of the light chains of cAC10. The Tub-tag ensures site-specific homogeneous conjugation and unique biophysical and physicochemical properties of TUB-010. The stable linkage between the highly potent payload MMAE to the antibody may be a crucial requirement for ADCs to be both effective and safe. Using long term *in vitro* incubation of TUB-010 or Adcetris in rat, cynomolgus or human sera, we demonstrated significantly higher stability of TUB-010 as compared to Adcetris. Consistent with these findings, *in vivo* experiments showed high amounts of linker-payload transferred to serum proteins in Adcetris-treated rats, whereas only negligible amount of linker-payload was transferred to serum proteins in TUB-010-treated rats. Observed differences in stability between the two ADCs can likely be attributed to the distinct conjugation technologies: maleimide-based for Adcetris and Tub-tag for TUB-010. ADCs that utilize maleimide-based technology for conjugation are susceptible to elimination of the maleimide through retro-Michael reaction which results in premature loss of linker-payload from the ADC (30, 31). Linker-payload is then transferred to serum proteins such as albumin and may be transported into multiple healthy tissues and cause off-target toxicities. Importantly, TUB-010 showed excellent *in vivo* stability in cynomolgus monkeys and rats with almost antibody-like *in vivo* pharmacodynamic data and no evidence of increased clearance supporting the superiority of Tub-tag technology over the maleimide-based technology. It should be noted that the chemical bond that is formed between 3-formyl-*L*-tyrosine and HA-VC-PAB-MMAE is an oxime. Even though oximes have been described to undergo hydrolysis, the remarkable stability of TUB-010 can be explained chemically by the proximity to the *p*-hydroxyl group of the tyrosine derivative, enabling a stabilization of the corresponding oximes via intramolecular hydrogen bonding (37, 38).

In addition to premature release of the cytotoxic drug, the non-specific uptake of the intact ADC into the target-negative healthy tissues may also contribute to undesirable toxicities. The non-specific uptake of ADCs is generally influenced by their physicochemical properties with positive charge, higher hydrophobicity, and a tendency to aggregate leading to increased unspecific cellular uptake (39). VC-PAB-MMAE is highly hydrophobic, and the hydrophobicity increases with the number of linker-payload moieties attached to the antibody. Unlike Adcetris, which is a heterogenous DAR 0-8 ADC, TUB-010 is a homogenous DAR 2 ADC. In addition, the Tub-tag is both the source of hydrophilicity which counterbalances the hydrophobic linker-payload, and a carrier of a negative charge. These unique properties of TUB-010 resulted in >5 fold reduced HMWS formation during storage at elevated temperatures as compared to Adcetris. Most importantly, these unique physicochemical properties of TUB-010 resulted in 2-10-fold lower cellular toxicity mediated by unspecific ADC uptake as compared to Adcetris in a panel of 10 target-negative primary human cell lines.

TUB-010 showed potent *in vitro* cytotoxicity against a panel of CD30+ cell lines representing established preclinical models for various human CD30+ lymphomas. The cytotoxic effect correlated with the level of CD30 expression as well as with the sensitivity of the tested cell lines to MMAE. When compared to Adcetris, TUB-010 showed similar *in vitro* cytotoxicity in equal MMAE concentrations (i.e. twice higher ADC concentrations), which is not surprising given that the cytotoxic effect was assessed in a static cell culture environment. Notably, when *in vivo* efficacy of TUB-010 vs Adcetris was compared in the Karpas-299 xenograft model, TUB-010 was similarly effective in equal ADC concentrations, despite delivering only half MMAE concentrations. Additionally, TUB-010 was more effective when compared at equal MMAE concentrations, likely reflecting its improved PK properties.

The improved biophysical and physicochemical properties of TUB-010 vs Adcetris discussed above translated also into significantly better tolerability of TUB-010 vs Adcetris in *in vivo* studies with rats and cynomolgus monkeys, the latter based on a comparison with published Adcetris data. In toxicity studies in cynomolgus monkeys, TUB-010 was very well tolerated and could be administered at doses up to 15 mg/kg without reaching the MTD and with no observed mortality. This contrasts with historical studies of Adcetris, where the MTD was 3 mg/kg and lethal dose was 6 mg/kg (36). Notably, hematological toxicity particularly neutropenia was markedly reduced in TUB-010-treated rats and cynomolgus monkeys compared to Adcetris-treated animals based on current (rats) or published data (cynomolgus monkeys) (36).

The therapeutic window, i.e. the dose range of a drug that provides effective therapy with minimal adverse effects, is a crucial parameter for estimating the clinical potential of TUB-010 in the treatment of CD30-positive hematological malignancies. Adcetris has a narrow therapeutic window, meaning that its effective dose is very close to its MTD. Importantly, the *in vivo* efficacy and safety data presented in the current study indicate that TUB-010 has a markedly widened therapeutic window as compared to Adcetris when dosed at the same ADC (5-fold) or MMAE (2.5-fold) concentrations. As a result, TUB-010 has the potential to be administered at higher doses and potentially for longer durations which may result in higher efficacy.

In conclusion, TUB-010 is a novel, next-generation CD30-targeting ADC which utilizes a similar linker-payload and mAb as Adcetris. However, TUB-010 uses a cutting-edge technology to stably attach the linker-payload to the mAb via a short hydrophilic linker, the Tub-tag. Unlike Adcetris, TUB-010 is a site-specific, homogeneous ADC, characterized by increased stability and hydrophilicity. Reflecting its improved design, preclinical data demonstrate comparable efficacy when TUB-010 and Adcetris are administered at equal ADC concentrations, and significantly higher efficacy when dosed at equal MMAE concentrations. Moreover, TUB-010 exhibits lower toxicity with at least 5-fold increase in the MTD compared to Adcetris. In summary, TUB-010 is expected to offer higher efficacy and lower toxicity in clinical setting and has a potential to improve clinical outcome of patients who undergo CD30-targeting therapy.

## Authors’ Disclosures

The authors declare competing financial interests: The technology described in the manuscript is part of pending patent applications by M.G., M.-A. K., D.S., H.L., C.P.R.H. and J.H. The following authors are employees of Tubulis GmbH: M.G., S.S., P.C., M.-A. K., I. M., G.F.-R., A. M. V., D.S. and J.H.

## Authors’ Contributions

**M. Gerlach:** Data curation, formal analysis, investigation, supervision, methodology, visualization, writing – review & editing. **S. Schmitt:** Data curation, formal analysis, investigation, supervision, methodology, visualization, writing – original draft. **P. Cyprys:** Investigation. **M.-A. Kasper**: investigation, methodology. **I. Mai:** Data curation, investigation. **M. Klanova:** writing – original draft.

**A. Maiser:** Investigation. **H. Leonhardt**: Supervision, project administration. **C.P.R. Hackenberger:** supervision, project administration. **G. Fingerle-Rowson:** Conceptualization, supervision, methodology, writing – review & editing. **A.M. Vogl:** Conceptualization, supervision, methodology, writing – original draft. **D. Schumacher:** Supervision, funding acquisition, methodology, project administration, writing – review & editing. **J. Helma:** Conceptualization, supervision, funding acquisition, methodology, writing – review & editing.

## Acknowledgements

This work was supported by grants from the German Federal Ministry for Economic Affairs and Energy and the European Social Fund with grants (to D. Schumacher and J. Helma; EXIST FT I); and the Bavarian Ministry of sEconomic Affairs, Regional Development and Energy with grants (to D. Schumacher, J. Helma; m4-Award).

TUB-010 is being developed by Oncoteq AG, Switzerland, under the company code TEQ102, through an exclusive license from Tubulis GmbH.

## Data availability statement

The data generated in these studies are available upon reasonable request from the corresponding author.

