## Supplement for "TUB-010, a novel anti-CD30 antibody-drug conjugate based on Tub-tag technology, widens the therapeutic window by reducing toxicity while maintaining high efficacy"

**Affiliations:**

### Supplementary Methods

#### Cell culture

Cell lines were purchased from the "German Collection of Microorganisms and Cell Cultures" (DSMZ, Leibniz Institute, Germany), Japanese Collection of Research Bioresources (JCRB) via Tebubio (France) or Merck (Germany): Karpas-299 (Merck, RRID:CVCL_1324), HL-60 (DSMZ, ACC 3, RRID:CVCL_0002), SR-786 (ACC 369, DSMZ), L-540 (ACC 72, DSMZ, RRID:CVCL_1362), L-428 (ACC 197, DSMZ, RRID:CVCL_1361), SU-DHL-1 (ACC 356, DSMZ, RRID:CVCL_0538), HDLM-2 (ACC 17, DSMZ RRID:CVCL_0009), DL-40 (JCRB1337-P, Tebubio, RRID:CVCL_2889), MTA (IFO50513-P, Tebubio, RRID:CVCL_3032), MOTN-1 (ACC 559, DSMZ, RRID:CVCL_2127), HH (ACC 707, DSMZ, RRID:CVCL_1280), BXPC-3 (ACC 760, DSMZ, RRID:CVCL_0186) and SW-620 (300466, Tebubio, RRID:CVCL_0547). Cells were cultured according to the manufacturer's instructions in RPMI 1640 medium supplemented with GlutaMAX (Gibco, Thermo Fisher Scientific, USA) and 10% or 20% fetal bovine serum (FBS; Gibco, Thermo Fisher Scientific, USA). The ExpiCHO-S cell line (Thermo Fisher, USA, RRID:CVCL_5J31) was cultured in ExpiCHO Expression Medium (Thermo Fisher Scientific, USA), the Expi293F cell line (Thermo Fisher Scientific, USA, RRID:CVCL_D615) in Expi293 Expression Medium (Thermo Fisher Scientific, USA). Healthy primary human cells of different origins were purchased from Innoprot (Spain) and cultivated in their proprietary media (Innoprot, Spain).

#### Antibody expression and purification

To produce cAC10 with Tub-Tag on the C-terminus of the light chain (cAC10 LC Tub), mammalian expression vectors pcDNA3.4-TOPO including the coding sequences for heavy and light chain IgG1 were ordered from Thermo Fisher Scientific (USA). The antibody was produced over 8 days in ExpiCHO-S or Expi293F cells (Thermo Fisher Scientific, USA) according to the manufacturer's instructions using a heavy to light chain DNA ratio of 1:1. After expression, 0.01% sodium azide, 4 mM EDTA, 1 µM pepstatin A, 10 µM leupeptin, 1.54 µM aprotinin (Carl Roth), were added to the supernatants and pH was adjusted to 7.2. For purification, protein A affinity chromatography (HiTRAP Fibro™ PrismA, Cytiva, USA) was performed on an Äkta pure (Cytiva, USA). The pH of the eluate was adjusted to 7.0-7.4 with 0.5 M NaOH followed by a buffer exchange into storage buffer using Amicon Ultra-4 centrifugal filter (Sigma-Aldrich, USA).

#### TTL expression and purification

Recombinant tubulin tyrosine ligase (TTL) was produced based on published protocols (1, 2). Briefly, the enzyme was expressed as recombinant Sumo-TTL fusion protein with an N-terminal $His_6$-Tag in *E. coli* BL21 (DE3) (New England Biolabs, USA). His-SUMO-TTL was purified using immobilized metal affinity chromatography (IMAC). Protein aliquots were snap-frozen and stored at -80 °C.

#### Chemicals

Chemicals were purchased from Carl Roth (Germany), Merck (Germany) or Thermo Fisher Scientific (USA) unless stated otherwise.

#### Analytical characterization

##### Analytical size exclusion chromatography (SEC)

Analysis via size-exclusion chromatography (HPLC-SEC) of TUB-010 was conducted on a Vanquish Flex (U)HPLC System using a MAbPac SEC-1 300 Å, 4 x 300 mm column (Thermo Fisher Scientific, USA) with a flow rate of 0.15 mL/min. Separation of different ADC/antibody (mAb) populations was achieved during a 30-minute isocratic elution using a phosphate buffer at pH 7 (20 mM $Na_2HPO_4$/$NaH_2PO_4$, 300 mM NaCl, 5% v/v isopropyl alcohol as a mobile phase. 8 µg ADC/mAb were injected. Quantification of monomer and HMWS was achieved by integration of the peak area at 220 nm.

##### Analytical hydrophobic interaction chromatography (HIC)

The (U)HPLC-HIC measurements were conducted on a Vanquish Flex UHPLC System with a MabPac HIC Butyl 4.6 x 100 mm column (Thermo Fischer Scientific, USA). Separation of different

ADCs/antibodies have been achieved with the following gradient: A: 1 M $(NH_4)_2SO_4$, 500 mM NaCl, 100 mM $NaH_2PO_4$ pH 7.4; B: 20 mM NaH2PO4, 20% (v/v) isopropyl alcohol, pH 7.4. 0% B: 0-1 min, 0-95% B: 1-15 min, 95% B: 15-12 20 min, 95-0% B: 20-23 min, 0% B: 23-25 min, with a flow of 700 µL/min. 12 µg sample were injected onto the column. UV chromatograms were evaluated at 220 nm.

**Liquid chromatography–mass spectrometry (LC-MS)**

Antibodies and ADCs were diluted to 0.2 mg/mL in PBS followed by reduction (5 mM DTT) and deglycosylation with PNGase F (0.5 U/mg, Promega, USA) at 37°C for 2 hours. The subsequent analysis was carried out with a LC-MS system consisting of the following core components: Acquity H-Class plus, Acquity TUV detector and Xevo G2-XS QTof (Waters, USA). A Acquity UPLC BEH300 C4 2.1 x 50 mm column (Waters) and the following gradient and flow rates were applied for desalting of the samples (A= $H_2O$ + 0.1% FA, B = ACN + 0.1% FA): 25% B, 0.2 mL/min, 0-1 min; 95%B, 0.2 mL/min, 1-3.5 min; 95%B, 0.4 mL/min, 3.5-4.5 min; 25%, 0.4 mL/min, 4.5-5 min; 95%B, 0.4 mL/min, 5-5.5 min; 25%B, 0.4 mL/min, 5.5-7.5 mL/min.

Samples were ionized in positive ion mode applying a capillary voltage of 1.50 kV and a cone voltage of 80 V with the analyser set in sensitivity mode. Source temperature was 120°C and desolvation temperature was 450°C. Cone gas flow was 50 L/h and desolvation gas flow was 850 L/h. Ionisation type was set to ESI with a mass range from 500 m/z to 3000 m/z. Raw data was analyzed with UNIFI.

#### Pharmacokinetics in rat

PK studies were conducted at Charles River Laboratories Edinburgh Ltd. in accordance with local authorities. Female Sprague Dawley rats (Crl:CD(SD), Charles River UK Limited; RRID:RGD_734476) were given a single intravenous injection at a dose level of 5 mg/kg of either TUB-010 or Adcetris (9 animals/group).

Blood was collected at 30 min, 1 h, 4 h, 24 h, 48 h, 96 h, 168 h, 336 h and 504 h (n=3 for each sampling point) with a target volume of 1 mL and processed to serum. Samples were analyzed for intact ADC and total antibody (mAb) as described below. Free MMAE concentrations were quantified via LC-MS. Subsequently pharmacokinetic parameters (e.g., $t_{max}$, $C_{max}$, $C_{max}$/dose, $T_{last}$, $t_{1/2}$, $AUC_{inf}$, $AUC_{tlast}$) were estimated using Phoenix WinNonlin pharmacokinetic software (Version 6.4, Certara, USA). A non-compartmental approach consistent with the intravenous bolus route of administration was chosen.

Transferred MMAE from the ADC to serum proteins at 24 and 48 h post treatment was analyzed by a LC-MS-based method as described previously (3).

#### Toxicology and toxicokinetics in rats

Toxicokinetic studies were conducted at Charles River Laboratories Hungary Kft. Female and Male Sprague Dawley rats (6-8 weeks old at onset of treatment) were dosed on Day 1, 8, 15, 22 at 10 mg/kg of either TUB-010 or Adcetris. Animals were observed at least twice daily, and body weights were recorded daily. Blood was collected from all animals for haematology, coagulation, and clinical chemistry on day 26 and from recovery animals on day 51. Samples for toxicokinetic analyses were collected on day 1, 8, 15 and 22 (pre- and post-dose) and analyzed using the same methods described for the pharmacokinetic analyses in rats. Gross pathology including macroscopic examination, bone marrow smears, organ weights and tissue preservation was performed on all animals.

**Analysis of total mAb and intact ADC by ELISA**

To evaluate the PK of the ADCs *in vivo*, the total antibody concentration was measured at different time points in serum of ADC-treated Sprague Dawley rats. Total antibody was analyzed in serum over the range 2000 – 15.6 ng/mL. Nunc 96-well plates (Thermo Fisher Scientific, USA) were coated with recombinant human CD30/TNFRSF8 Protein (R&D systems, USA) and sealed with PCR foil. Plates were incubated in a fridge to maintain a temperature between 2-8°C overnight. The coated plates were washed 3x with PBST (PBS + 0.05% Tween, Merck, Germany), blocking solution (2 % albumin in PBST, Carl-Roth, Germany) was added and the plate was incubated at room temperature for 1 hour. The coated plates were washed 3x with PBST. Standards, quality controls (QCs) and test samples were added, the plates were sealed and incubated at room temperature for 1 hour. The plates were washed 3x with PBST. Anti-human IgG (γ-chain specific)-peroxidase antibody (Sigma-Aldrich, USA; Cat# A8419,

RRID:AB_258388) was added and incubated for 1 hour at room temperature. The plates were washed 3x PBST. TMB (Thermo Fisher Scientific, USA) was added, the plates were sealed and incubated at room temperature for 10 minutes. 1 M sulfuric acid (Carl-Roth, Germany) was added. Using a microplate reader Infinite 200 PRO (Tecan, Switzerland), the absorbance at a wavelength of 450 nm was measured.
To evaluate the stability of the ADCs *in vivo*, the intact ADC concentration was measured at different time points in serum of ADC-treated Sprague Dawley rats. Intact ADC was analyzed in rat serum over the range 2000 – 15.6 ng/mL. Nunc 96-well plates were coated with rabbit anti-VC-PAB-MMAE pAb (Levena Biopharma, USA; Cat# LEV-PAE1-100) and sealed with PCR foil. Plates were incubated in a fridge to maintain a temperature between 2-8°C overnight. The coated plates were washed 3x PBST, blocking solution (2 % albumin in PBST) was added, the plate was sealed and incubated at room temperature for 1 hour. The coated plates were washed 3x PBST. Standards, QCs and test samples were added, the plates were sealed and incubated at room temperature for 1 hour. The plates were washed 3x PBST. HRP-labeled goat anti-human IgG (H+L) (Abcam, UK; Cat# ab209700) was added and incubated for 1 hour at room temperature. The plates were washed 3x PBST. TMB was added, the plates were sealed and incubated at room temperature for 10 min. 1 M sulfuric acid was added. Using a microplate reader Infinite 200 PRO (Tecan, Switzerland), the absorbance at a wavelength of 450 nm was measured.

#### Cell binding

To determine equilibrium binding constants ($K_D$; as an avidity measurement), CD30-positive cells were incubated with antibodies at increasing concentrations up to saturation and stained with an αhuman IgG heavy and light chain secondary antibody (Thermo Fisher Scientific, USA; Cat# A-11013, RRID:AB_141360). Data points were normalized to the maximum mean fluorescence intensity (MFI) and analyzed by a non-linear regression using a one-site specific binding model. To determine maximum binding, CD30-positive cells were incubated with the antibodies at a concentration of 5 µg/mL followed by secondary antibody staining and MFI ratios were calculated by dividing the specific MFI by the secondary antibody background control.

#### Determination of antibodies bound per cell

Antibodies bound per cell (ABCs) were determined by using the QIFIKIT (Agilent, USA). For this, cells were stained with 10 µg/mL of mouse anti-human CD30 (BioLegend, UK; Cat# 333902, RRID:AB_1134022) or an irrelevant mouse isotype control (BioLegend, UK; Cat# 400102, RRID:AB_2891079). The primary antibodies were detected by a fluorescein-conjugated anti-mouse secondary antibody. Cells were acquired by flow cytometry on a CytoFLEX LX flow cytometer (BD Biosciences, USA). Quantification of ABCs as an estimation for surface receptor expression was performed by simultaneous acquisition of secondary antibody labelled setup and calibration beads consisting of bead populations with known number of antigens on the surface to generate a standard curve. The fluorescence is correlated with the number of bound primary antibody molecules on the cells and on the beads.

#### ELISA-based binding

To determine binding of total αCD30 antibodies to soluble CD30, flat-bottom surface-treated 96-well-plates (Nunclon, Thermo Fisher Scientific, USA) were coated with recombinant human CD30 (R&D systems, USA). After blocking with TBST + 2 % bovine serum albumin (Carl Roth, Germany), varying concentrations of αCD30 antibodies or antibody-containing serum samples were allowed to bind. Bound proteins were detected by anti-human IgG (γ-chain specific)-AP antibody (Jackson Immunoresearch, USA; Cat# 309-055-008, RRID:AB_2339661). 4-Methylumbelliferyl phosphate substrate (Merck, Germany) was added and reaction was stopped by 3 M NaOH. Fluorescence was measured at excitation/emission of 385 nm/448 nm on a microplate reader Infinite M1000 Pro (Tecan, Switzerland).

#### Internalization by flow cytometry

For pHrodo-based investigation of internalization, antibodies were labeled with pHrodo™ Deep Red Antibody Labeling Kit (Thermo Fisher Scientific, USA) according to manufacturer's instructions. CD30-positive and -negative cell lines were incubated with 5 µg/mL pHrodo Deep Red-antibodies for 1 h, 5 h and 24 h at 37°C. An increase in MFI indicates the presence of αCD30 antibodies in late endosomal and lysosomal compartments. The MFI ratio was determined by deviding the MFI of pHrodo-incubated cells by the MFI of unstained cells.

#### Internalization by structured illumination microscopy

To assess internalization of antibodies by CD30-positive cell lines, brentuximab derivatives and ADCs were labeled with Alexa Fluor™ 594 Antibody Labeling Kit (Thermo Fisher Scientific, USA) according to the manufacturer's instructions. CD30-positive cells were incubated for 24 h at 37°C or for 1 h at 4°C with 5 µg/mL labeled antibodies. For samples incubated at 4°C, membrane staining with an AlexaFluor 488-labeled αHLA-DR antibody (BioLegend, UK; Cat# 307620, RRID:AB_493175) was performed. Samples incubated at 37°C were fixed and permeabilized using the Cytofix/Cytoperm Fixation/Permeabilization Kit (BD Biosciences, USA) according to the manufacturer's instructions and late endosomes and lysosomes were stained using αLAMP1-AlexaFluor 488 (Thermo Fisher Scientific, USA; Cat# 53-1079-73, RRID:AB_657534). For all conditions, post-fixation took place with 4% paraformaldehyde followed by nuclear staining with 1 µg/mL of DAPI (Thermo Fisher Scientific, USA). Next, cells were transferred onto coverslips using a Shandon Cytospin 3 cytocentrifuge (Thermo Fisher Scientific, USA). Finally, the coverslips were mounted onto glass slides using Vectashield (Vectorlabs, USA).

3D structured illumination microscopy (SIM) acquisition was done on a Deltavision OMX V3 microscope (General Electric, USA) equipped with a 100 × 1.4 oil immersion objective UPlanSApo (Olympus, Japan), 405, 488, and 593 nm diode lasers and Cascade II EMCCD cameras (Photometrics, USA). Raw data were first reconstructed and corrected for color shifts with the provided software softWoRx 6.0 Beta 19 (unreleased). A custom-made macro in Fiji finalized the channel alignment and established composite TIFF stacks.

#### Characterization of Fc effector functions

##### ADCC

For the Calcein release-based antibody-dependent cellular cytotoxicity (ADCC) assay, peripheral blood mononuclear cells (PBMCs) were isolated from healthy donor buffy coats (purchased from DONAS GmbH, Germany) using LeucoSep tubes (Greiner Bio-One, Austria). For this, 15 mL of Histopaque®-1077 (density 1.077 g/mL, Merck, Germany) was added to LeucoSep tubes and centrifuged for 1 min at 1000 g to move the Histopaque solution below the insert. Then, the buffy coats are mixed 1:1 with DPBS (Thermo Fisher Scientific, USA) and added carefully to the LeucoSep tubes (max. 35 mL per tube). Tubes were centrifuged for 10 min at 1000 g without break. The upper part of plasma is removed and PBMC fraction is collected, washed with DPBS and counted. Natural killer (NK) cells were then MACS-sorted from PBMCs using human NK cell isolation kit (Miltenyi Biotec, Germany) by negative selection according to the manufacturer's instructions yielding untouched human primary NK cells.

CD30-positive target cells were stained with 16 µM Calcein AM (Thermo Fisher Scientific, USA). NK and target cells were then incubated at an effector-to-target ratio of 3:1 for 4 h at 37°C in presence of increasing concentrations of cAC10, cAC10 LC, TUB-010, Adcetris, αMHC-I (Anti-HLA Class I, Invivogen, USA) and a commercial human IgG1 isotype (BioLegend, UK; Cat# 403501, RRID:AB_2927629) up to saturation (100 nM). Cells permeabilized with 2.5% Triton X (Sigma-Aldrich) served as positive control. Supernatants were transferred to a flat black non-binding 96-well plate (Greiner Bio-One, Austria) and fluorescence was measured at 485/535 nM via the Infinite M1000 Pro reader (Tecan, Switzerland).

The percent specific killing was calculated by dividing the Calcein released by antibody-mediated killing minus background Calcein release (NKs + targets) from Calcein released by Triton X-permeabilized cells (maximum killing) minus background Calcein release (targets only). We determined either maximum ADCC/specific killing at 15 µg/mL or generated a concentration-dependent killing curve that was analyzed by a non-linear regression using a one-site specific binding model.

#### ADCP

For the Calcein stain-based antibody-dependent cellular phagocytosis (ADCP) assay, PBMCs from healthy donors were generated as described before. Monocytes were then MACS (magnetic cell separation)-sorted using human classical monocyte isolation kit (Miltenyi Biotec, Germany) by negative selection according to the manufacturer's instructions yielding untouched human primary monocytes. Monocytes were seeded on flat-bottom surface-treated 96-well-plates (Nunclon, Thermo Fisher Scientific, USA) in RPMI 1640 + 10% FBS containing 100 ng/mL M-CSF (Peprotech, USA) and differentiated for 7 days into M2-like macrophages. CD30-positive target cells were stained with 5 µg/mL Calcein red-orange (Thermo Fisher Scientific, USA) and macrophages (effectors) with 0.5 µg/mL Calcein AM (Thermo Fisher Scientific, USA). To setup the antibody-dependent cellular phagocytosis (ADCP) assay, target cells and adherent macrophages were incubated at an effector-to-target ratio of 1:1 for 3 h in presence of increasing concentrations of cAC10, cAC10 LC, TUB-010, Adcetris, αMHC-I (Anti-HLA Class I, Invivogen, USA) and a commercial IgG1 isotype (BioLegend, UK) up to saturation (100 nM). At the end of the incubation time, non-adherent target cells were harvested and the adherent macrophages were detached by a DPBS solution (Thermo Fisher Scientific, USA) containing 4 mg/mL lidocain-HCl (Sigma-Aldrich, USA) and 5 mM EDTA (Sigma-Aldrich, USA). Afterwards, cells were acquired by flow cytometry. The percentage of specific phagocytosis was analyzed by determining the percentage of Calcein AM- and Calcein red-orange-positive cells from macrophages.

#### CDC

Complement-dependent cytotoxicity (CDC) was measured by incubating CD30-positive cells with cAC10, cAC10 LC, TUB-010, Adcetris, αMHC-I (Anti-HLA Class I, Invivogen, USA) and an IgG1 isotype for 2 hours in presence of RPMI 1640 and 20% human serum (HS) of two healthy donors as sources of complement or of commercial heat-inactivated (h.i.) HS (Sigma-Aldrich, USA). Cells were stained with Fixable Aqua Dead Cell Stain Kit (Thermo Fisher Scientific, USA) and analyzed by flow cytometry to determine the percentage of viable cells.

#### DC maturation assay and immunogenic cell death

Monocytes from PBMCs were isolated as described before. Monocytes were seeded in flat-bottom 24-well plates with surface treatment for maximum adhesion (Nunc, Thermo Fisher Scientific, USA) and cultured for 72 h in RPMI 1640 + 1.5% human serum (HS, human male AB plasma, Sigma-Aldrich) supplemented with 800 IU/mL of GM-CSF (Peprotech, USA) and 580 IU/mL of IL-4 (Peprotech, USA) to generate immature monocyte-derived DCs (iDCs). At the same time, L-540 cells were pre-incubated with MMAE, TUB-010, Adcetris, αCD30-MMAF normalized to a total toxin concentration of 1 µM for 72 h. Afterwards, supernatant of pre-incubated L-540 cells containing free toxins was transferred to iDCs a final dilution of 1:1 and incubated for another 24 h. As a positive control, 1 µg/mL LPS (Thermo Fisher Scientific, USA) was added to iDCs. After 24h, cells were stained with CD86-APC (BioLegend, UK; Cat# 374207, RRID:AB_2721448) and Fixable Aqua Dead Cell Stain Kit and analyzed by flow cytometry. As markers for immunogenic cell death, Calreticulin positive cells, HMGB1 and ATP release were quantified. For all readouts, CD30-positive cells were incubated with 100 nM TUB-010 and MMAE for 24 h – 72 h. Calreticulin staining of cells was performed using PE anti-Calreticulin antibody (Abcam, UK) or respective isotype control. Lumit™ HMGB1 Human/Mouse Immunoassay (Promega, USA) and RealTime-Glo™ Extracellular ATP Assay (Promega, USA) was used to quantify secreted HMGB1 and ATP into the media according to manufacturer's instructions.

#### Flow cytometry

Flow cytometry measurements were performed on a CytoFLEX LX flow cytometer (Beckman Coulter, USA) with FlowJo 10.6 (Tree Star Inc., USA).

#### Data analysis

Calculations were performed using Excel 365 (Microsoft, USA) and graphs were generated using GraphPad Prism 9 (GraphPad, USA).

### Supplementary Figures, Tables and Results

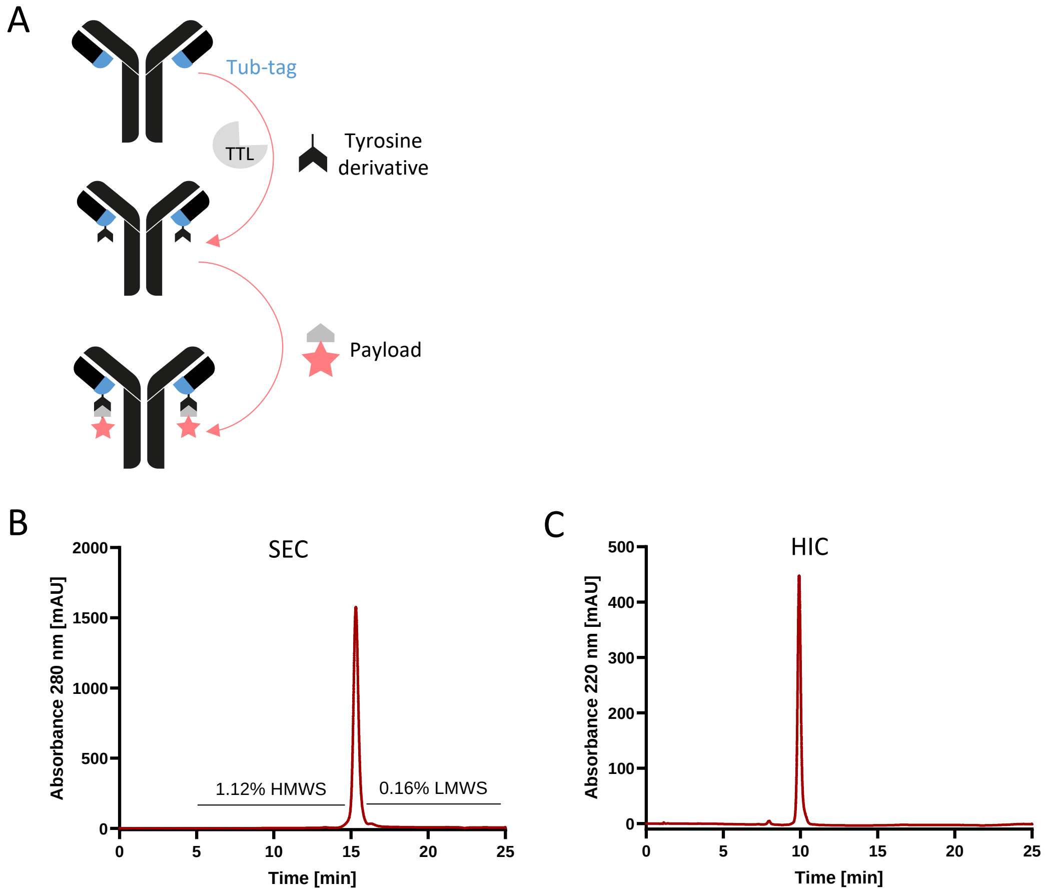


**Figure S1. TUB-010 is conjugated to the payload MMAE via Tub-Tag technology and represents a highly homogenous and stable ADC**
**(A)** Schematic overview of the Tub-tag conjugation process. **(B-C)** Analysis of TUB-010 by HPLC-SEC **(B)** and HPLC-HIC **(C).** HMWS: high molecular weight species, LMWS: low molecular weight species.

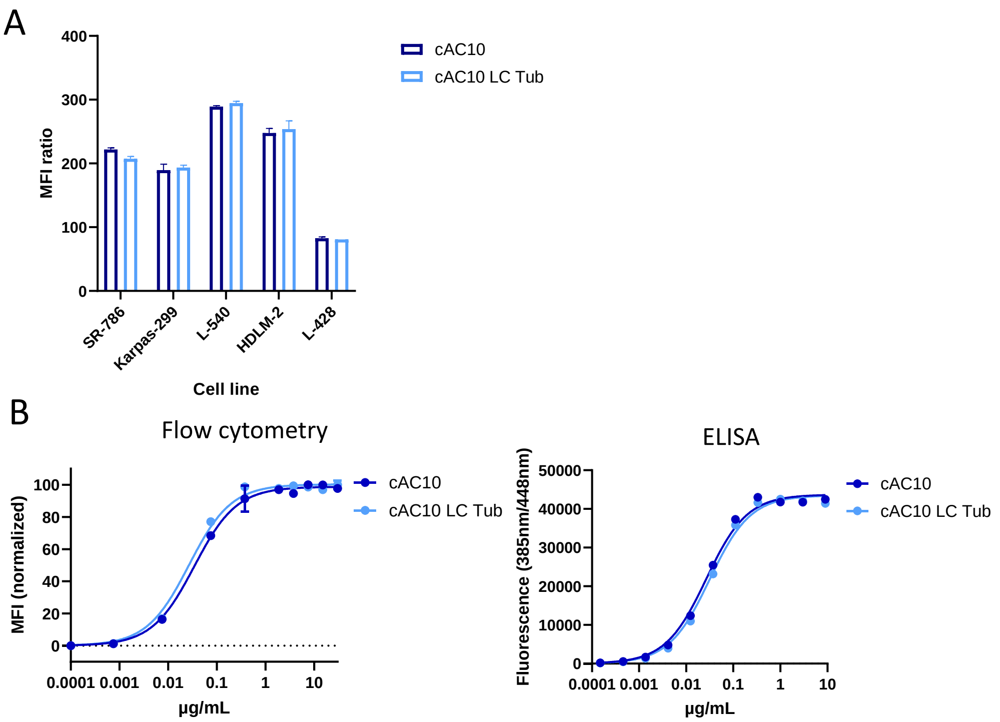


| Molecule | $K_D$ Flow cytometry | $K_D$ ELISA |
|---|---|---|
| cAC10 | 0.034 µg/mL (0.23 nM) | 0.026 µg/mL (0.18 nM) |
| cAC10 LC Tub | 0.027 µg/mL (0.17 nM) | 0.031 µg/mL (0.21 nM) |

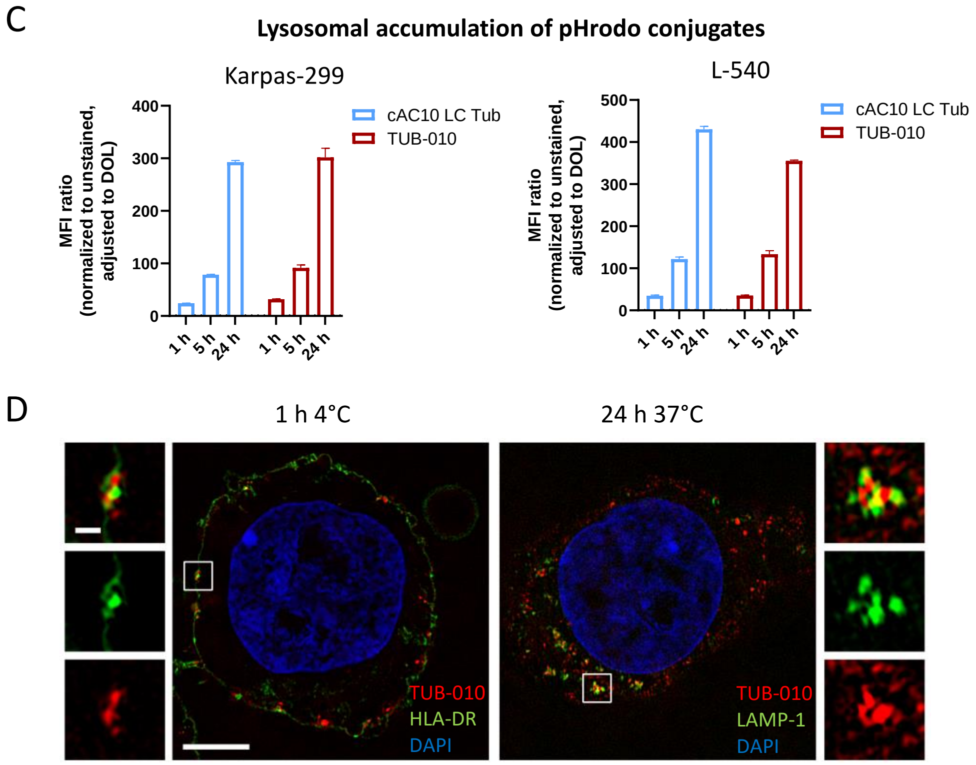


**Figure S2. Brentuximab and derivatives show high affinity binding to CD30 and strongly internalize into CD30-positive cells** **(A)** Maximum binding of cAC10 and cAC10 LC Tub at 5 µg/mL to CD30-positive cell lines measured by flow cytometry. Graph shows mean of n = 2 ± SEM. **(B)** Binding curves of cAC10 and cAC10 LC Tub on Karpas-299 cells and to soluble recombinant human CD30 determined by flow cytometry and ELISA, respectively. Graph shows mean of n = 2 ± SEM. For flow cytometry, MFIs were normalized to 100% for all molecules to eliminate any differences in maximum binding. Calculated $K_D$ values are shown in table. **(C)** Karpas-299 and L-540 cells were incubated for indicated time points with saturating concentrations of pHrodo Deep Red-labeled cAC10 LC Tub and TUB-010 and analyzed by flow cytometry. MFI ratio was determined by dividing the fluorescence of antibody-treated cells from unstained cells and normalized to the degree of labeling (DOL) of the antibody conjugates. Graph shows mean of n = 2 ± SEM. **(D)** Representative images of 3D structured illumination microscopy

(SIM) to visualize TUB-010 internalization. L-540 cells were incubated for 24 h at 37°C or for 1 h at 4°C (control) with 5 µg/mL of AF594-labeled TUB-010. At 4°C co-staining of the membrane with αHLA-DR-AF488 was performed. At 37°C the lysosomes were visualized by αLAMP1-AF488. All conditions were counterstained with DAPI. Images were adjusted to saturation for better visibility of internalization using Image J. The scale bar represents 5 µm and in the enlarged images 0.5 µm.

A

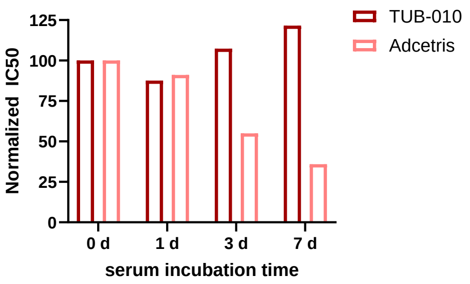


B

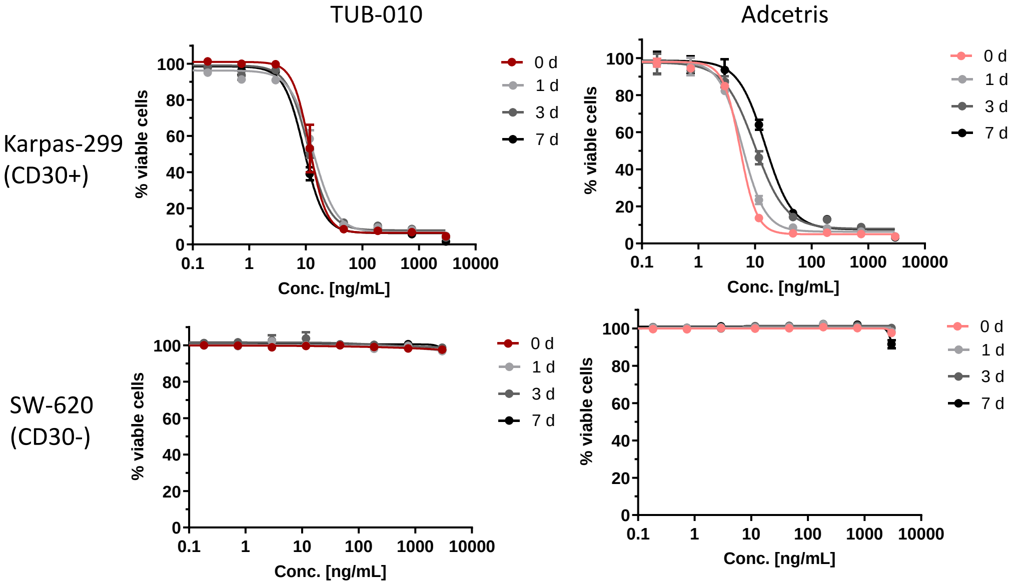


C

Study design:

| | |
|---|---|
| Rats | Sprague Dawley Crl:CD(SD) |
| Group size | *n* = 9 (females) |
| Administration route | i.v. (tail vein) |
| Administration | Single dose |
| Dose level | 5 mg/kg TUB-010 and Adcetris |

D

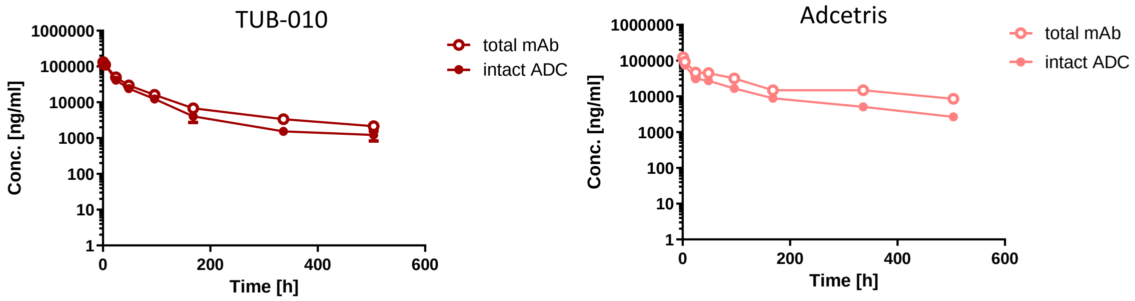


E

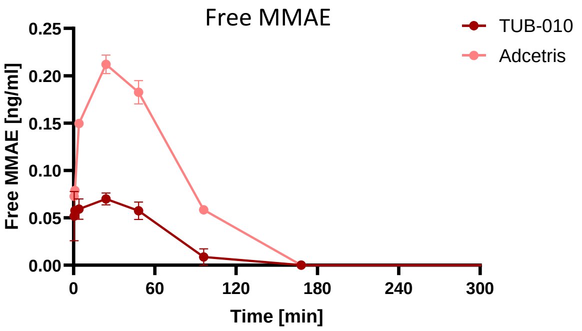

**Figure S3. TUB-010 exhibits higher *in vitro* and *in vivo* serum stability compared to Adcetris**

**(A-B)** TUB-010 and Adcetris were incubated for 0-7 days in human serum at 37°C and afterwards applied in a cytotoxicity assay on CD30-positive Karpas-299 and CD30-negative SW-620 cells. IC50 values **(A)** and killing curves **(B)** of the resazurin-based cytotoxicity readout are shown. Data points represent mean ± SEM, n = 2. (**C-E)** Pharmacokinetic analysis of TUB-010 and Adcetris in rats. (**C)** Study design. **(D)** Total antibody and intact ADC were measured in rat serum samples by ELISA after indicated sampling times. Total ADC was captured using an anti-idiotype brentuximab antibody, intact ADC by an anti-MMAE antibody. Data points represent mean ± SEM, n = 3. **(E)** Free MMAE was measured in serum samples by LC-MS. Data points represent mean ± SEM, n = 3.

A

| Indication | Cell line | CD30 level | Antigen density per cell | MMAE sensitivity (IC50) [nM] |
|---|---|---|---|---|
| **ALCL (ALK+)** | SR-786 | high | 325,146 | 0.07 |
| | Karpas-299 | med | 213,390 | 0.03 |
| | SU-DHL-1 | low | 118,750 | 0.24 |
| **HL** | L-540 | high | 440,332 | 0.20 |
| | HDLM-2 | med | 291,197 | 0.03 |
| | L-428 | low | 112,273 | 1.52 (upregulation of MDR1 confirmed) |
| **ALCL (ALK-)** | DL-40 | high | 340,744 | 0.09 |
| **CTCL** | HH | high | 437,538 | 0.14 |
| **PTCL** | MTA | high | 394,559 | 0.17 |
| | MOTN-1 | high | 393,745 | 0.08 |

B

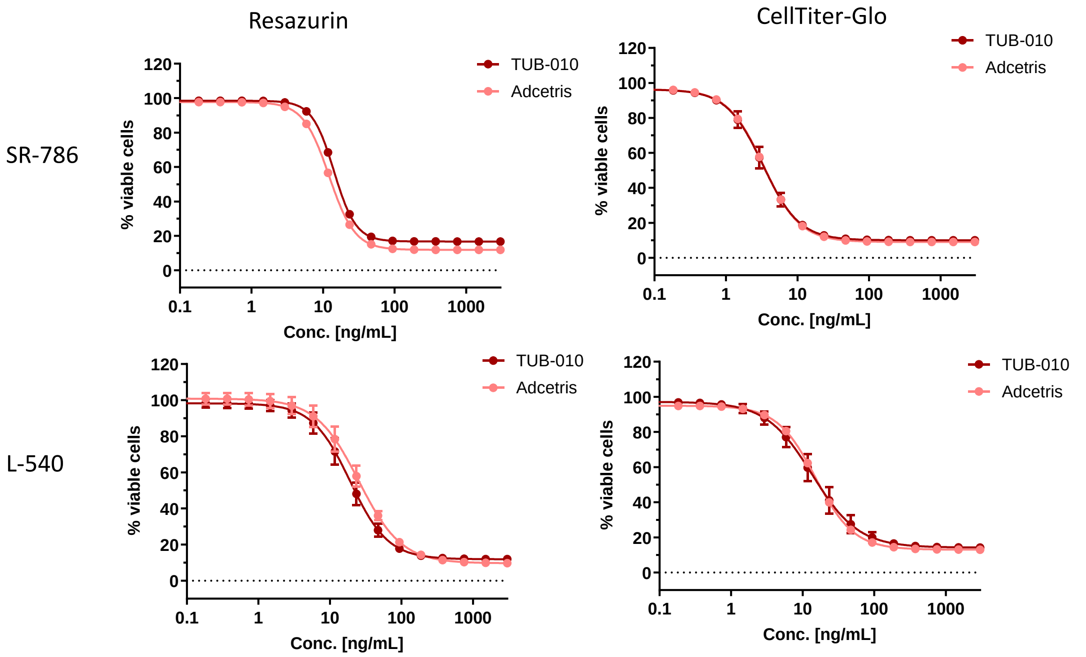


C

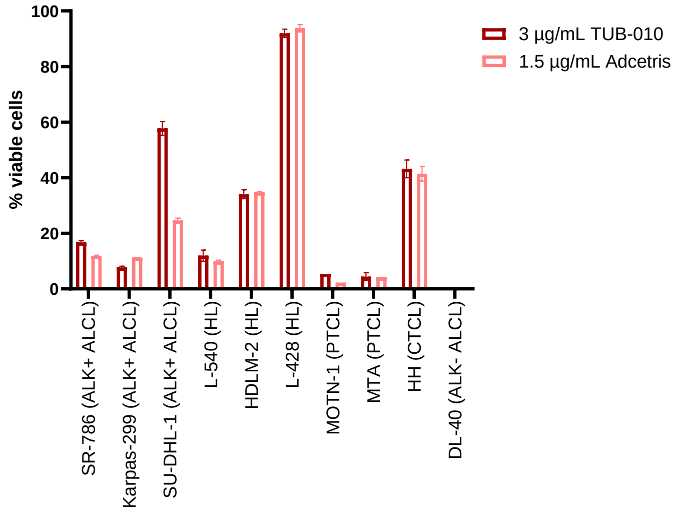


**Figure S4.** ***In vitro*** **cytotoxicity of TUB-010 and Adcetris**
**(A)** Antigen density and MMAE sensitivity of cancer cell lines from different indications. **(B)** ADC-mediated concentration-dependent cytotoxicity on $CD30^{high}$ cell lines SR-786 (ALK+ ALCL) and L-540 (HL) analyzed by resazurin- and CellTiter Glo-based killing readout. ADC concentrations were adjusted for MMAE amounts. Indicated concentrations are valid for TUB-010, while half of the concentration of Adcetris was used. **(C)** Cytotoxicity of TUB-010 versus Adcetris at indicated concentrations. ADC concentrations were adjusted for MMAE amounts. Data points represent mean ± SEM, n = 2.

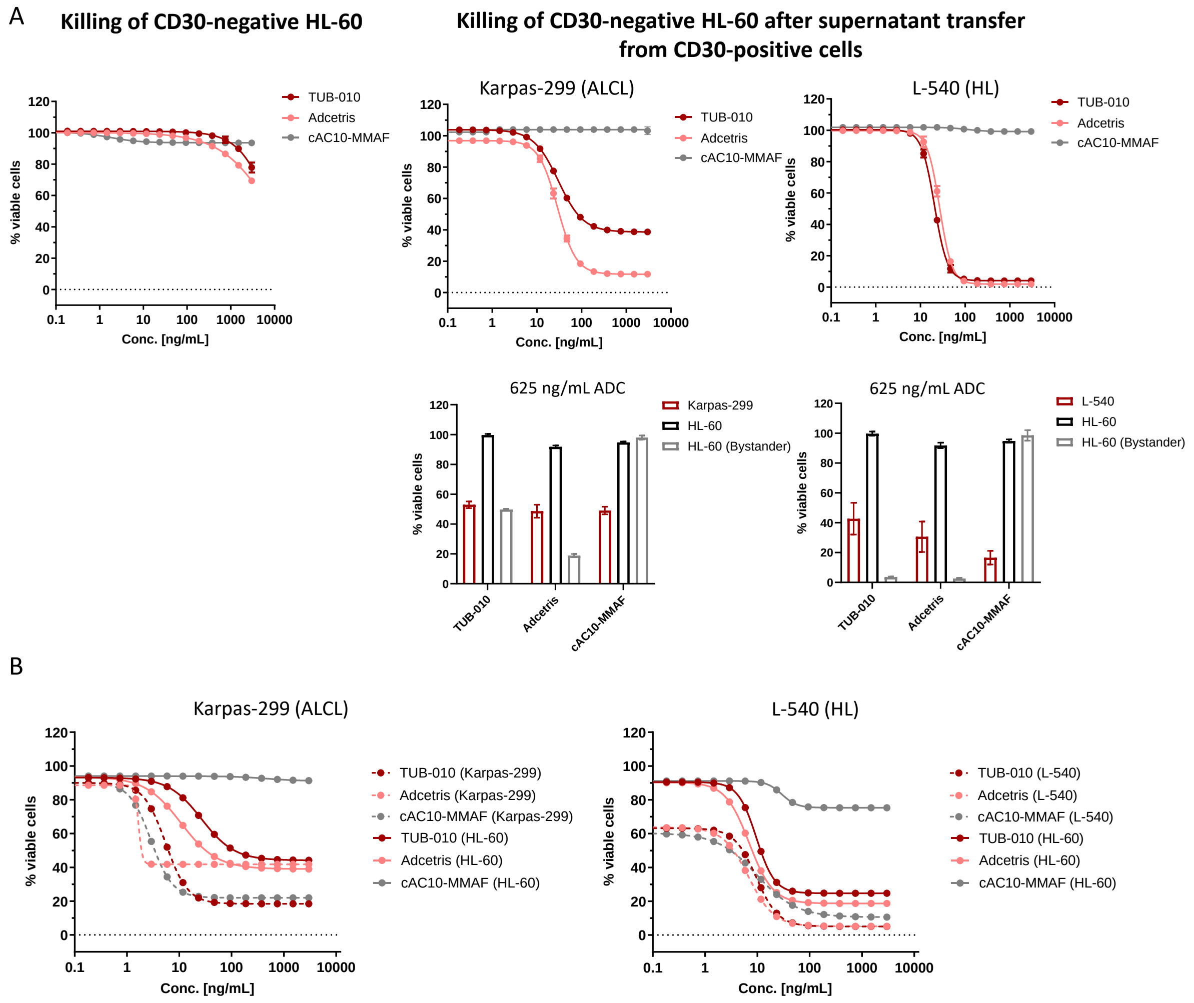


**Figure S5. TUB-010 and Adcetris exhibit bystander activity *in vitro***

**(A)** Supernatant from TUB-010- and Adcetris-preincubated CD30-positive cells (Karpas-299 and L-540) was transferred to CD30-negative HL-60 cells and killing was analyzed by a resazurin-based viability readout. ADC concentrations were adjusted for MMAE amounts. Indicated concentrations are valid for TUB-010, while half of the concentration of Adcetris was used. Bar graphs show direct and indirect cytotoxicity of TUB-010 versus Adcetris at indicated concentrations on CD30-positive and -negative cells. Data points represent mean of duplicates ± SEM. **(B)** To investigate bystander activity in co-culture, CD30-positive and -negative cells were co-cultured in presence of TUB-010 and Adcetris. In a flow cytometry-based killing readout including a live/dead stain, dead CD30-positive and -negative cells could be distinguished.

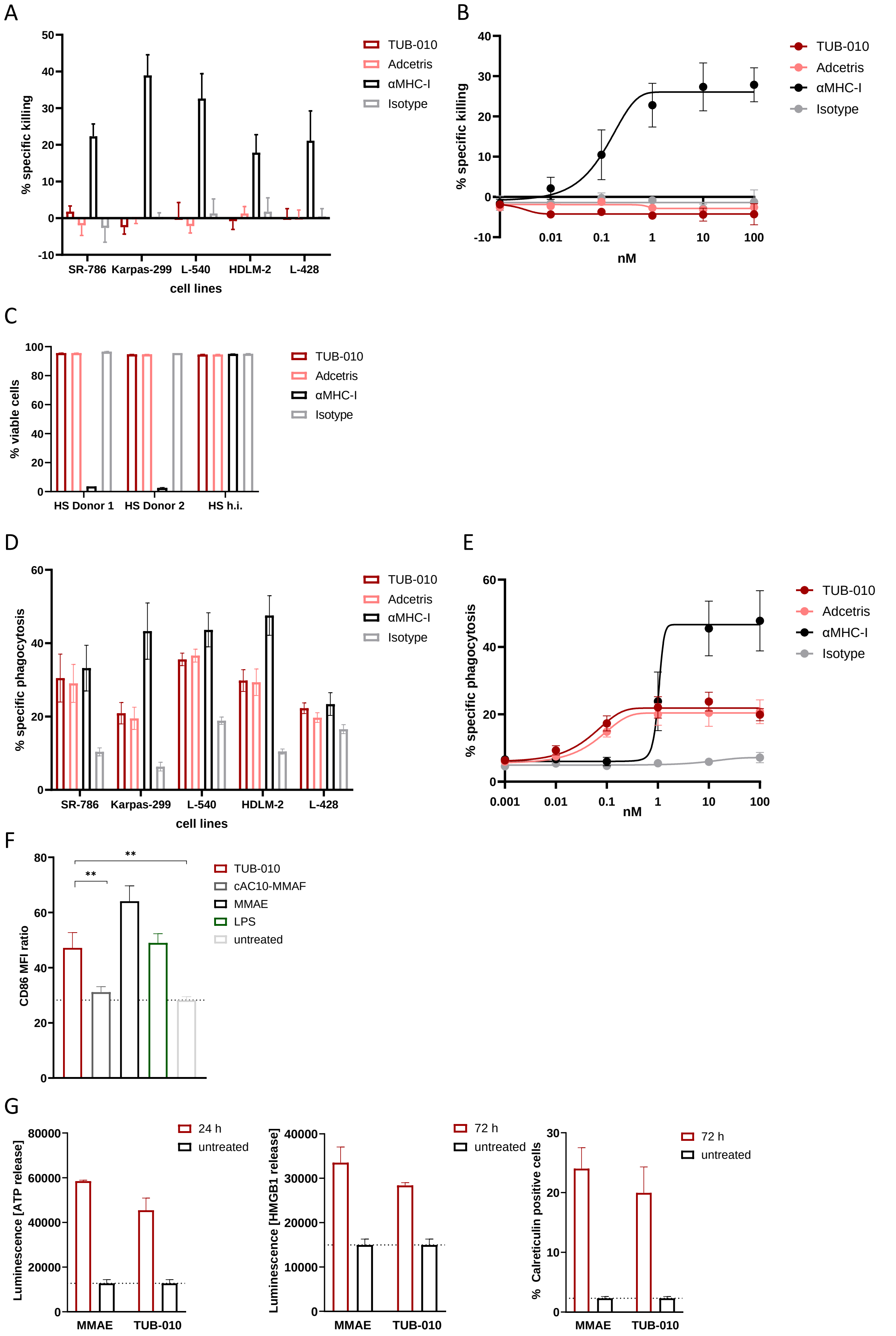


**Figure S6. TUB-010 exhibits ADCP, but not ADCC and CDC activity and is induces markers of immunogenic cell death**

**(A-B)** Calcein release-based antibody-dependent cellular cytotoxicity (ADCC) readout. Co-cultures of healthy donor (HD)-derived NK cells and Calcein-stained CD30-positive tumor cells were incubated with 15 µg/mL **(A)** or increasing concentrations **(B)** of indicated mAbs and ADCs. % specific killing was calculated by dividing the Calcein released by antibody-mediated killing from Calcein released by Triton X-permeabilized cells (maximum killing). Bars show means of n = 6 healthy donor NK cells with SEM as error bars. Concentration-dependent killing was evaluated on Karpas-299 cells using n = 3 HD NK cells. (**C)** Complement-dependent cytotoxicity (CDC) was measured by incubating CD30-positive cells with indicated mAbs and ADCs in presence of medium and 20% human serum (HS) of two healthy donors (HS donor 1 and 2) as sources of complement or of commercial heat-inactivated (h.i.) HS. Cells were stained with a live dead stain and analyzed by flow cytometry to determine the percentage of viable cells. Bars show means of duplicates with SEM as error bars. **(D-E)** Antibody-dependent cellular phagocytosis (ADCP) was measured by incubating primary Calcein AM-stained HD-derived macrophages with Calcein red-orange-stained CD30-positive cell lines in presence of 15 µg/mL **(D)** or increasing concentrations **(E)** of mAbs and ADCs. % specific phagocytosis was analyzed by determining the percentage of Calcein AM- and Calcein red-orange-positive cells from macrophages. Bars show means of n = 6 healthy donor macrophages with SEM as error bars. Concentration-dependent killing was evaluated on Karpas-299 cells using n = 3 HD macrophages. **(F)** DC maturation. Supernatant from L-540 cells, pre-incubated with Adcetris, TUB-010, αCD30-MMAF or MMAE, normalized to 1 µM MMAE/MMAF, was transferred to iDCs to investigate CD86 upregulation by flow cytometry as a readout for DC maturation. As a positive control, iDCs were maturated with LPS. The graphs show means of n = 8 different donors with SEM as error bars. For statistical analysis, a Wilcoxon-signed rank test was applied. **(G)** Immunogenic cell death induced by TUB-010 and MMAE (100 nM) on L-540 cells. As markers for immunogenic cell death, ATP release and HMGB1 release were measured by a luminescence-based assay and Calreticulin exposure on the cell membrane was measured by flow cytometry. Data points represent means ± SEM.
